# Small aberrant viral genomes induce the innate immune response to arenaviruses

**DOI:** 10.64898/2026.03.09.710519

**Authors:** Abraham Ayanwale, Wanda Christ, Léna Vandenabeele, Silke Olschewski-Pawlita, Saskia Johanns, Chris Hoffmann, Lisa Oestereich, Maria Rosenthal, Thomas Pietschmann, Benjamin E. Nilsson-Payant

## Abstract

Arenaviruses are a family of negative-sense RNA viruses mainly found in rodents in which they cause chronic asymptomatic infections. Some arenaviruses, however, can occasionally infect humans and cause pathogenic viral haemorrhagic fevers. These viral haemorrhagic fevers are associated with a massively dysregulated immune response characterized by excessive release of pro-inflammatory cytokines resulting in hyperinflammatory immunopathology. Here, we systematically characterize the immune response of human cells to a wide panel of mammalian arenaviruses and identify a clear distinction between New World viruses, which uniformly induce a robust interferon response, and Old World viruses, which do not induce any significant host response. This innate immune response is primarily driven by RIG-I-mediated RNA sensing and might be aided by differences in viral interferon antagonist expression patterns. Furthermore, we identified differences in quantity and type of non-standard viral genomes produced in unpassaged Old World and New World virus infections. Finally, we demonstrate that small non-standard viral RNA expression by New World viruses are key drivers of the innate immune response and might help explain how the host response can shape viral disease phenotypes and inform triage of patients.

## Introduction

The *Mammarenavirus* genus in the *Arenaviridae* family (henceforth called arenaviruses) are a highly diverse group of negative-sense bisegmented RNA viruses with an ambisense genome within the class of the *Bunyaviricetes.* They comprise some of the most pathogenic human viruses, including Lassa virus (LASV) and Junín virus (JUNV)^1,2^. Arenaviruses typically circulate in rodents or bats and – depending on their geographical distribution and genetic phylogeny – can be classified into Old World arenaviruses (OWA), mainly found in Africa and Eurasia, and New World arenaviruses (NWA), which circulate in the Americas ^3^. Each virus is typically associated with a single specific natural host species, in which they cause asymptomatic and often persistent infections and transmit the virus through the oral-faecal route. Some arenaviruses, however, are known to occasionally cause zoonotic infections and human disease. Human infections are primarily thought to occur through the inhalation or ingestion of animal secretions or contaminated dust particles, but human-to-human transmission has been observed for LASV and JUNV through direct contact with bodily secretions of viraemic patients and through nosocomial infections. Arenavirus-induced haemorrhagic fevers are associated with flu-like symptoms, gastrointestinal and neurological symptoms, which in severe cases can culminate in internal and external vascular haemorrhaging, systemic shock, and multi-organ failure resulting in death. Fatal cases of haemorrhagic viral fevers usually exhibit systemic viral dissemination and cytokine storms which account for the severe immunopathology. Currently, there are no approved vaccines with the exception of the live-attenuated Candid#1 JUNV vaccine and treatment options are limited to symptoms-based supportive care and the broadly antiviral ribavirin.

It has long been known that RNA viruses produce defective viral genomes (DVGs) or non-standard viral genomes (nsVGs). This has been attributed to the fact that the viral RNA-dependent RNA polymerase (RdRp) is error-prone and relies on a multitude of complex cellular interactions to maintain faithful replication of viral genomes. Traditionally it has been assumed that the generation of nsVGs had an attenuating or interfering influence on viral infections due to several reasons: Firstly, packaging of these nsVGs results in defective interfering particles (DIPs) that cannot produce infectious progeny virus without the presence of complementing helper viruses. Secondly, nsVGs tend to be shorter than their respective wildtype genomes and are therefore replicated more quickly and thus serve as a sink for viral and cellular resources, preventing resource allocation to full genome replication. Finally, nsVGs have been suggested to be highly immunogenic, thus activating the antiviral host cell defences^4^. However, recently the accumulation of different classes of nsVGs have been directly associated with increased disease severity and viral pathogenicity for several highly inflammatory viruses, including influenza A virus (IAV), respiratory syncytial virus (RSV) and SARS-CoV-2^5–9^. Interestingly, arenavirus nsVGs have previously been described, identifying the formation of several distinct nsVG species^10–13^. Nevertheless, the impact of abundance and type of nsVG on viral pathogenicity remains unknown.

In this study, we aimed to understand the molecular mechanisms underlying innate immune activation of different arenaviruses and how nsVGs contribute to these processes. We observed that NWA induce a robust antiviral host response in different relevant human cell types, while OWA only induce a very limited or delayed transcriptional response. This differential immune response was not correlated to the ability of different viruses to inhibit the antiviral response. Induction of the type I and II interferon (IFN) response was directly linked to sensing by RIG-I and signalling through MAVS, while PKR was also shown to play an IFN-independent critical role in controlling virus infections. Both NWA and OWA were able to produce nsVGs but exhibited distinct differences in abundance and type that could account for the different observed antiviral host response phenotypes.

## Results

### NWA, but not OWA induce a robust antiviral host response in human lung cells

To compare the antiviral host response in relevant human cells to different arenaviruses, we infected adenocarcinomic human alveolar basal epithelial A549 cells – commonly used to study virus-host interactions of respiratory viruses including arenaviruses – with a wide range of pathogenic and apathogenic NWA (highlighted in red colours throughout this study) and OWA (highlighted in blue colours throughout this study), spanning all relevant clades of mammarenaviruses (Fig. 1a). Since arenaviruses were propagated in African green monkey cells in which they induce secretion of cytokines including IFN, we purified virus stocks from soluble small proteins before virus infections to prevent virus-independent cross-signalling of monkey cytokines in human cells.

**Figure 1:**
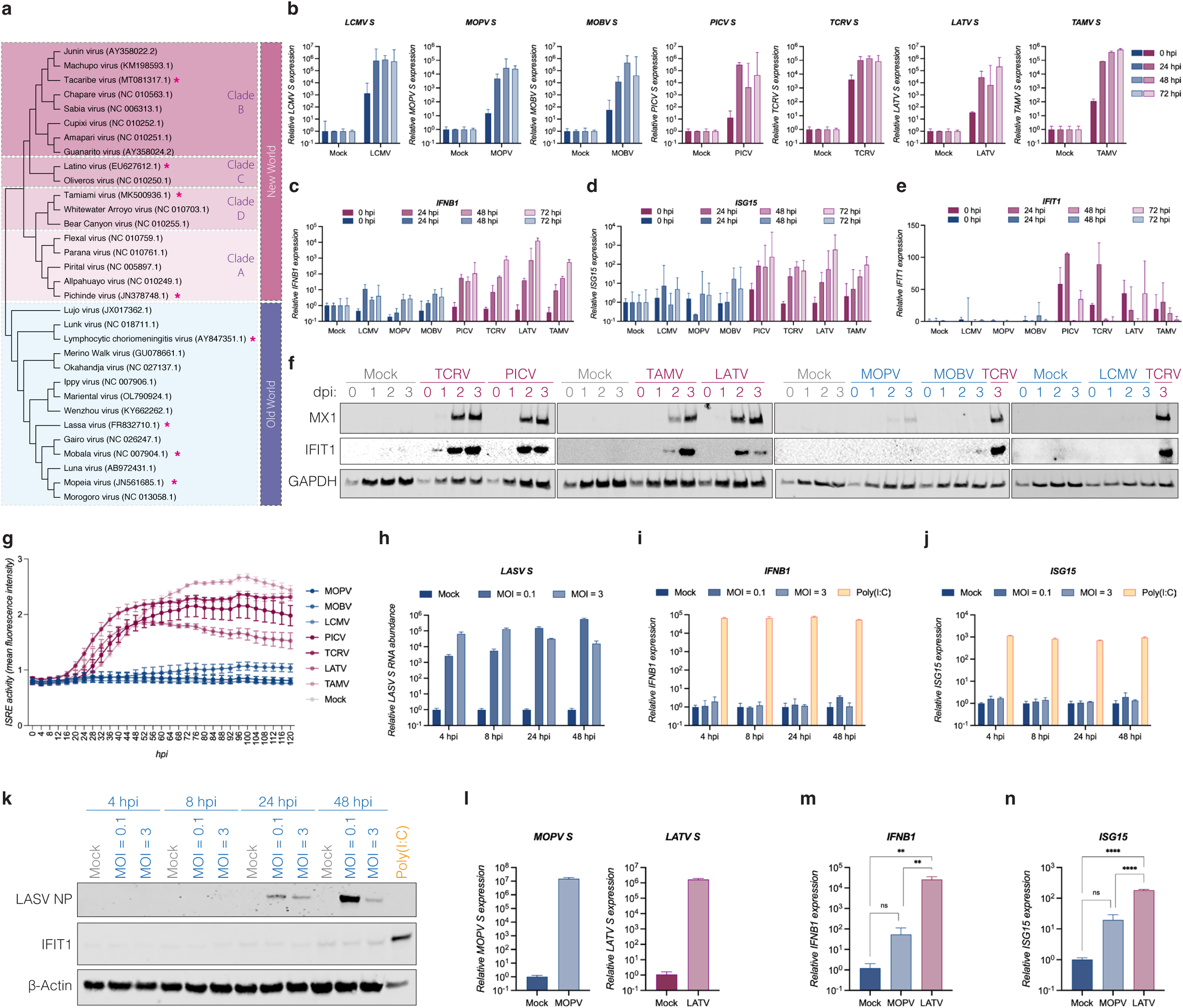
NWA induce a robust type I IFN response. (a) Maximum-likelihood phylogenetic tree based on the amino acid sequences of arenavirus RdRps. Arenaviruses are grouped into Old World and New World (clades A-D) viruses. Viruses used in this study are highlighted with a red asterisk (*). Old World arenaviruses are highlighted in blue colours; New World arenaviruses are highlighted in red colours throughout this study. (b-f) A549 cells were infected with the indicated arenaviruses at an MOI of 0.1 and samples were collected immediately after infection (0 hpi) or over a 72 h timecourse and analysed by RT-qPCR for the expression of (b) viral S segment RNA, (c) *IFNB1*, (d) *ISG15* or (e) *IFIT1* gene expression or by (f) western blot analysis for Mx1, IFIT1 and GAPDH protein expression. (g) A549-ISRE-mScarlet cells were infected with the indicated arenaviruses at an MOI of 0.1 and ISRE-dependent reporter activity was measured at 4 h intervals for 120 hours post-infection (hpi). The graph depicts the mean fluorescence intensity from three independent biological replicates with error bars representing the standard deviation of the mean. (h-k) A549 cells were infected with LASV at an MOI of 0.1 or 3 and RNA or whole cell lysates were extracted at the indicated time points. A549 cells transfected with poly(I:C) RNA for 24 h were used as control. Extracted RNA was analysed by RT-qPCR for (h) LASV S segment, (i) *IFNB1* or (j) *ISG15* gene expression. (k) Whole cell lysates were analysed by western blot for the expression of LASV NP, IFIT1 or ϕ3-Actin. (l-n) HPMECs were infected with MOPV or LATV at an MOI of 0.1 for 48 hpi. Total RNA was analysed by RT-qPCR for (l) viral S segment expression, (m) *IFNB1* or (n) *ISG15* gene expression. All graphs representing RT-qPCR data depict the mean relative fold change of gene expression normalized to human GAPDH expression levels from three independent biological replications. Error bars represent the standard deviation of the mean. Statistical significance was determined by One-way ANOVA with post-hoc Tukey test for multiple comparisons: **** (*p* < 0.0001), *** (*p* < 0.001), ** (*p* < 0.01), * (*p* < 0.05), ns (*p* > 0.05).

A549 cells were infected at a low multiplicity of infection (MOI = 0.1) with the OWA Lymphocytic choriomeningitis virus (LCMV), Mobala virus (MOBV) and Mopeia virus (MOPV), as well as the NWA Pichinde virus (PICV), Tacaribe virus (TCRV), Latino virus (LATV) and Tamiami virus (TAMV), and compared replication dynamics and host cell responses over a 72 h timecourse. RT-qPCR analysis of the relative abundance of the viral S segment (NP gene) for each virus demonstrated that all viruses were able to infect and replicate in A549 cells, with a strong increase in viral RNA in the first 24 h of infection, followed by plateauing viral RNA levels (Fig. 1b). Despite these largely similar replication dynamics for all viruses, we observed strikingly different patterns of activation of the host cell antiviral responses. While *IFNB1* (Fig. 1c), *ISG15* (Fig. 1d) and *IFIT1* (Fig. 1e) gene expression was limited in A549 cells infected with OWA (LCMV, MOBV, MOPV), cells infected with NWA universally exhibited strong transcriptional activation of these genes. These findings were validated at the protein level, where robust IFIT1 and Mx1 protein expression – two typical IFN stimulated antiviral proteins – was observed by 48 hpi for TCRV, PICV, LATV and TAMV, but not for LCMV, MOPV and MOBV (Fig. 1f). These data were reproduced and validated in separate experiments (Extended Data Fig. 1a -d).

In order to acquire an even higher kinetic resolution of the type I/III IFN response, A549 cells expressing a fluorescent reporter (mScarlet) under the control of an IFN stimulated response element (ISRE) were infected with both OWA and NWA and reporter expression was measured at 4 h intervals for 5 days (Fig. 1g). Notably, IFN-mediated reporter activity could be observed for all NWA from 24 hpi onwards, with strong increases in reporter activity until 48 hpi, after which the signal plateaued or slowly decreased. In contrast, no or only very limited reporter activity could be detected in A549 cells infected with LCMV, MOBV or MOPV.

Since MOBV and MOPV are apathogenic and not associated with human infections, and LCMV largely is associated with subclinical or mild disease in healthy individuals, we next infected A549 cells with the highly pathogenic OWA Lassa virus (LASV) at both low (MOI = 0.1) and high (MOI = 3) MOI to account for potential initial virus-load differences in virus-host interactions. Interestingly, we observed similar patterns to the apathogenic OWA at both MOIs, where we see robust viral infection and replication (Fig. 1h and Extended Data Fig 1a), but neither induction of *IFNB1* (Fig. 1i and Extended Data Fig 1b) or *ISG15* (Fig. 1j and Extended Data Fig 1c) gene expression nor IFIT1 protein expression (Figs. 1k and Extended Data Fig. 1d). In order to rule out potentially reduced infection rates of human cells by OWA, we performed immunofluorescence imaging of A549 cells infected with MOBV (Extended Data Fig. 1e) or LASV (Extended Data Fig. 1f) and verified that the majority of cells were efficiently infected by both viruses.

To validate whether these findings can be reproduced in a different relevant and primary cell model, human pulmonary microvascular endothelial cells (HPMECs) – representing one of the cell types where primary arenavirus infections take place – were infected with MOPV or LATV as representative members of OWA and NWA, respectively. At 48 hpi, efficient viral RNA replication could be detected for both viruses (Fig. 1l), but despite limited induction of *IFNB1* (Fig. 1m) and *ISG15* (Fig. 1n) after MOPV infection, LATV induced a significantly higher antiviral host response.

In summary, these data clearly demonstrate that NWA, but not OWA induce a strong antiviral host immune response in relevant human pulmonary cells.

### TCRV infection induces a comprehensive transcriptional host response

In order to comprehensively characterize the host transcriptional response to arenavirus infections in human cells, we infected A549 cells with LASV (a highly pathogenic OWA) and TCRV (a close genetic relative of highly pathogenic NWA) at an MOI of 0.1 and performed total RNA sequencing (RNA-seq) of RNA extracted at 48 hpi. Sample analysis immediately revealed that the transcriptional landscape of LASV infected cells looked relatively similar to their uninfected controls, while TCRV infection significantly changed the cellular transcriptome compared to their uninfected controls (Extended Data Fig. 2a-b). We subsequently analysed differentially expressed genes (DEGs) in infected compared to uninfected cells and found that LASV infections hardly induced any significant transcriptional changes (adjusted *p*-value <0.05 and |log2FC| >2), while TCRV infection induced hundreds of DEGs, most of them being overexpressed in infected cells (Fig. 2a-b). These DEGs in TCRV infected cells were highly enriched for biological processes related to immune processes and apoptosis (Fig. 2c). Analysis of the gene expression of specific gene sets related to the interferon alpha response (Fig. 2d), interferon gamma response (Fig. 2e), inflammatory responses (Fig. 2f) and apoptosis (Fig. 2g) clearly highlights that TCRV strongly upregulates these pathways, while LASV largely remains transcriptionally inert.

**Figure 2:**
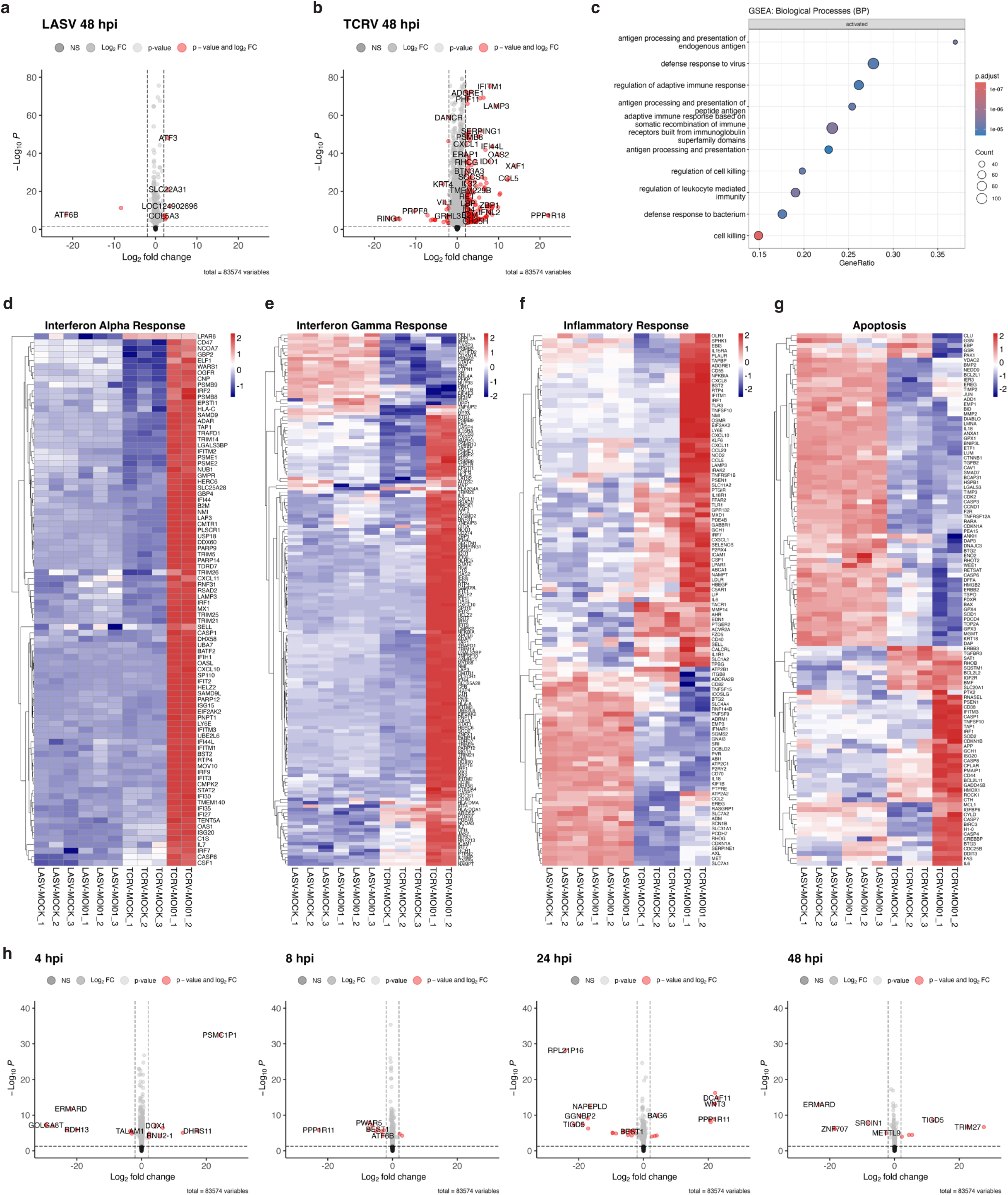
TCRV but not LASV induces comprehensive transcriptional changes in A549 cells. A549 cells were infected with TCRV or LASV at an MOI of 0.1 for 48 hpi and total RNA was analysed by total RNA-seq. (a-b) Volcano plots depict differentially expressed genes in (a) LASV infected and (b) TCRV infected cells compared to their respective uninfected control cells. (c) Gene set enrichment analysis of differentially expressed genes in TCRV infected cells compared to uninfected control cells. (d-g) Heatmaps depict gene expression levels of Hallmark gene sets related to (d) interferon alpha response, (e) interferon gamma response, (f) inflammation or (g) apoptosis. (h) A549 cells were infected with LASV at an MOI of 3 and total RNA was analysed by total RNA-seq at the indicated time points. Volcano plots depict differentially expressed genes in LASV infected cells compared to uninfected control cells at the same time.

In order to establish whether LASV induces any short-lived transient transcriptional responses which might be overlooked at a single time point, we infected A549 cells with LASV (MOI = 3) and performed RNA-seq at 4, 8, 24 and 48 hpi. However, we only observed very low levels of DEGs throughout the entire course of infection, starting at 4 hpi and with no discernible increase in DEGs over time (Fig. 2h and Extended Data Fig. 2c-d). Together, these data reveal that TCRV, but not LASV, induces significant transcriptional changes in infected cells and specifically upregulates pathways related to IFN signalling and apoptosis.

### Both NWA and OWA can effectively inhibit IFN induction, but not IFN signalling

We next sought to investigate the molecular basis of these distinct differences in host cell response to NWA and OWA, respectively. First, we established whether active viral replication is required to induce the observed IFN response to NWA. We infected A549 cells with equivalent amounts of UV inactivated and untreated TCRV and analysed IFIT1 protein expression and STAT1 phosphorylation as a measure for the type I/III IFN response (Fig. 3a). Here, we confirmed that the antiviral response is clearly induced by actively replicating virus and not by replication-incompetent viral particles or virus stock contaminants. Next, we infected A549 cells with LASV, MOPV or TCRV (MOI = 0.1) and extracted total RNA at 48 hpi and tested whether total and deproteinated RNA itself can induce an IFN response in naïve cells. Extracted RNA was transfected into A549-ISRE-Nluc2AGFP cells expressing interferon-inducible NanoLuc luciferase and reporter activity was measured 24 h post-transfection (Fig. 3b). RNA from uninfected cells did not induce any reporter activity, while RNA from cells infected with all three viruses induced comparable IFN responses. This indicates that the deproteinated viral RNA itself is sufficient to induce RNA sensing and that there are no inherently different RNA modifications impacting RNA sensing between OWA and NWA.

**Figure 3:**
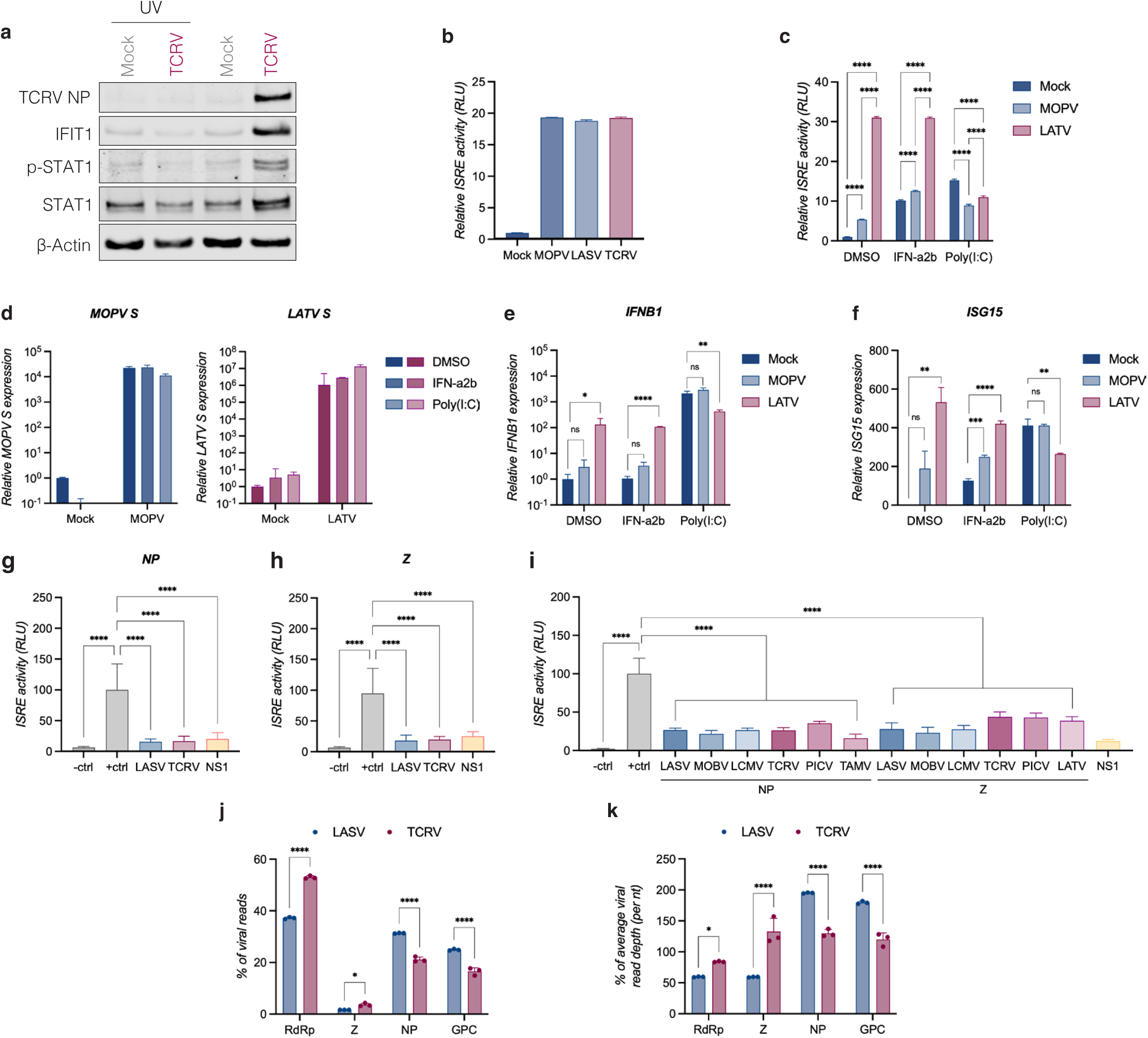
Both NWA and OWA can effectively antagonise RNA sensing and IFN induction. (a) A549 cells were infected with untreated or UV-irradiated TCRV at an MOI of 0.01 for 72 h. Whole cell lysates were analysed by western blot for TCRV NP, IFIT1, STAT1, phospho-STAT1 (Y701) expression. (b) Total RNA was extracted at 48 hpi from uninfected and MOPV, LASV or TCRV (MOI = 0.1) infected A549 cells. RNA was subsequently transfected into A549-ISRE-Nluc2AGFP cells and ISRE-induced NanoLuc luciferase activity was measured at 24 h post-transfection. The graph depicts the mean reporter activity normalised to uninfected control cellular RNA from six technical replicates. (c-f) A549-ISRE-Nluc2AGFP cells were infected with LATV or MOPV (MOI = 0.1) for 72 hpi before being treated with 1,000 iu/ml IFN-ɑ2b or 1 µg poly(I:C) RNA. ISRE-mediated NanoLuc luciferase activity was measured 24 h post-treatment. (c) The graph depicts mean relative light units normalised to the uninfected and untreated control from three independent biological replicates. Error bars represent the standard deviation of the mean. RT-qPCR analysis of (d) LATV and MOPV S RNA, (e) *IFNB1* and (f) *ISG15* gene expression levels normalised to the *ACTB* housekeeping gene. (g-i) HEK293T cells were transiently co-transfected with plasmids expressing a Firefly luciferase reporter under the control of an IFNB1 promoter, RIG-I and the indicated (g) NP or (h) Z proteins of LASV and TCRV, or (i) the NP and Z proteins of the indicated arenaviruses. The graphs depict the mean relative light units normalised to the uninhibited positive control from 6 to 10 technical replicates. Error bars represent the standard deviation of the mean. (j-k) A549 cells infected with TCRV or LASV (MOI = 0.1) for 48 hpi were analysed by total RNA-seq and viral ORF expression was determined relative to (j) the total number of viral reads or (k) to the size of ORFs and the mean viral reads per nucleotide. Statistical significance was determined by One-way ANOVA with post-hoc Turkey test (for e-f), One-way ANOVA with post-hoc Dunnett’s test (for g-i) and or by Two-way ANOVA with post-hoc Šídák’s test (for c, j-k) for multiple comparisons: **** (*p* < 0.0001), *** (*p* < 0.001), ** (*p* < 0.01), * (*p* < 0.05), ns (*p* > 0.05).

In order to establish whether any differences in host transcriptional responses could be explained by different abilities to antagonise IFN induction and signalling, we treated MOPV or LATV infected A549-ISRE-Nluc2AGFP cells (MOI = 0.1, 72 hpi) with recombinant IFN-ɑ2b or poly(I:C) RNA for 24 h and measured IFN-inducible reporter activity (Fig. 3c). In DMSO treated control cells we observed a similar phenotype as before with MOPV inducing low reporter activity and LATV inducing significantly stronger reporter levels. Treatment with IFN-ɑ2b resulted in a similar pattern, however, with increased reporter activity in uninfected and MOPV infected cells compared to DMSO treated cells. Finally, poly(I:C) transfections in uninfected cells induced a strong IFN response compared to DMSO treatment, but in both MOPV and LATV infected cells reporter activity was decreased compared to uninfected but poly(I:C) transfected cells. We validated these reporter assays by RT-qPCR for viral RNA expression (Fig. 3d) and endogenous levels of *IFNB1* (Fig. 3e) and *ISG15* (Fig. 3f) gene expression, confirming that neither MOPV nor LATV were able to inhibit IFN-ɑ2b-induced signaling. In contrast, LATV was able to strongly inhibit RNA sensing and poly(I:C) mediated IFN induction and MOPV, which exhibited low levels of IFN induction in untreated and IFN-ɑ2b-treated cells compared to uninfected cells, did not exhibit an increase in IFN response compared to uninfected cells. Since we were able to show that arenaviruses can antagonise RNA sensing, but not IFN sensing/signalling, we next investigated the IFN antagonistic potential of the viral NP and Z proteins – known IFN antagonists of arenaviruses – of TCRV and LASV outside of the context of viral infections. Here, we transiently overexpressed an IFN reporter together with either NP (Fig. 3g) or Z (Fig. 3h) proteins, using the potent IFN antagonist IAV NS1 protein as a positive control. Since we previously showed that the RNA sensing pathway could be inhibited by arenaviruses, we induced IFN induction by the overexpression of RIG-I. Surprisingly, we found that both NP (Fig. 3g) and Z (Fig. 3h) proteins efficiently antagonised RIG-I induced *IFNB1* transcription at a comparable level to IAV NS1 and that no significant differences could be observed between TCRV and LASV proteins. We then widely compared NP and Z proteins for different arenaviruses and observed that they all had comparable abilities to antagonise IFN induction when expressed in isolation (Fig. 3i). To address whether there are significant differences in expression levels of NP and Z between OWA and NWA, we quantified expression levels for all four arenavirus open reading frames (ORFs) from our RNA-seq data. Interestingly, we found that LASV had higher relative expression levels of the NP and GPC ORFs compared to RdRp and Z, whereas TCRV had a much more equal distribution of gene expression (Fig. 3j-k). These data suggest that LASV potentially expresses more NP compared to TCRV, while TCRV expressed higher levels of Z than LASV.

These data demonstrate that both OWA and NWA have the capacity to effectively inhibit RNA sensing and IFN activation, which is induced by active viral replication, but that potential inherent differences in viral protein expression levels might help shape the antiviral host response.

### TCRV generates higher levels of nsVGs than LASV

First characterisations of arenavirus nsVGs have recently been published^12,13^. However, the impact of arenavirus nsVGs on the immune response has not been investigated, yet. Therefore, we set out to characterize the aberrant replication landscape of LASV and TCRV in A549 cells and the frequency of nsVG generation in primary infections without serial passage of virus stocks. Here, we infected A549 cells with LASV or TCRV at an MOI of 0.1 for 48 h and identified nsVGs using next-generation sequencing using a custom-adapted bioinformatics pipeline based on the ViReMa algorithm: short-read Illumina sequencing can only identify nsVG with certainty if sequencing reads span a non-canonical junction, *i.e.* partly align to one region of the viral genome and partly align to another non-continuous region of the genome (Fig. 4a). Sequencing reads originating from nsVGs that do not span the junction site cannot be distinguished from reads stemming from full viral genomes. For both LASV (Extended Data Fig. 3a) and TCRV (Extended Data Fig. 3b) robust genome coverage of both viral segments was observed using reads correctly and fully aligned to their respective viral genome references. Interestingly, LASV genome coverage was markedly higher than that of TCRV and this was reflected in significantly more viral reads aligned to LASV (approximately 2.29% of human aligned reads) compared to TCRV (approximately 0.28% of human aligned reads) genomes (Fig. 4b).

**Figure 4:**
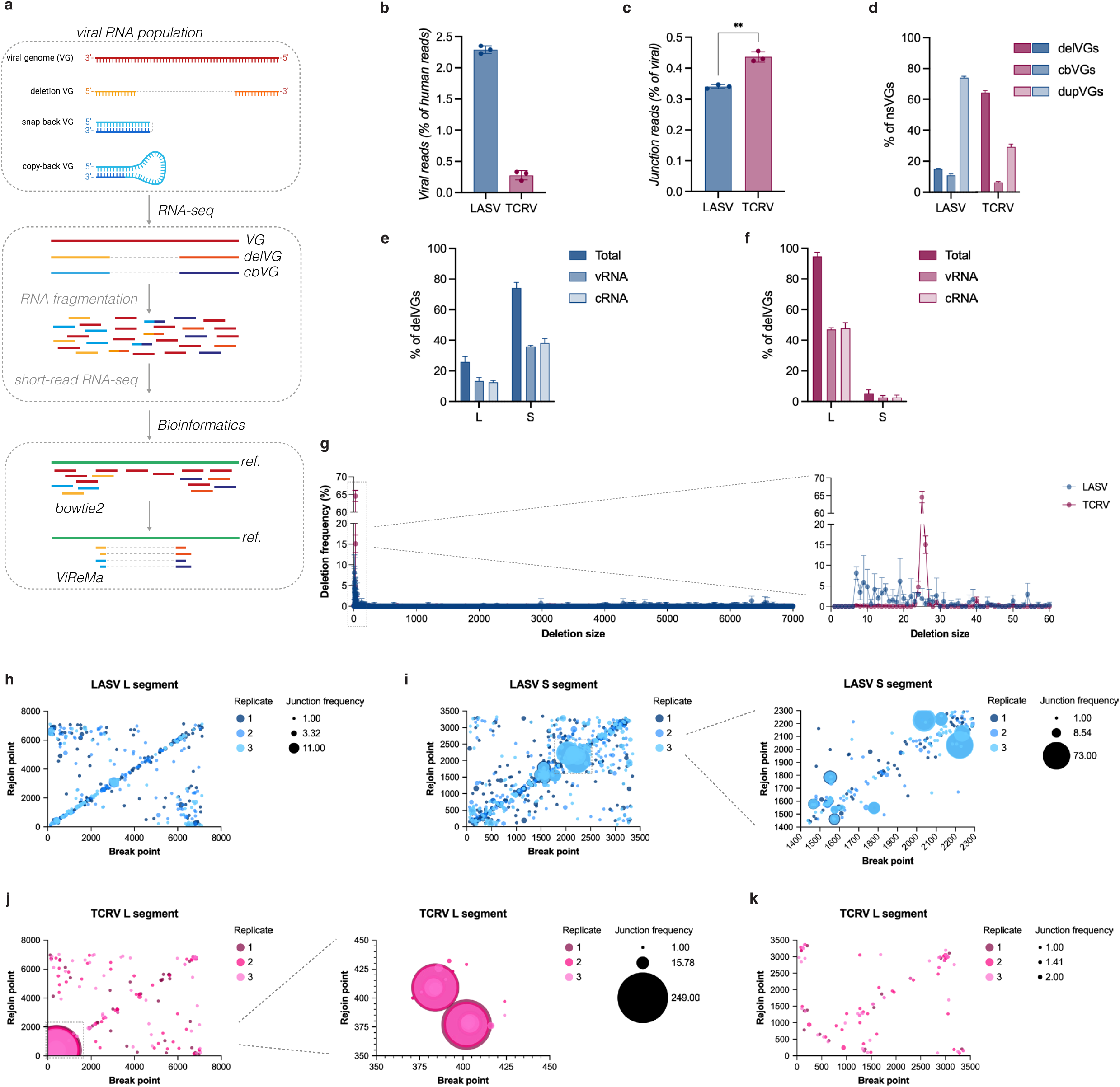
Analysis of TCRV and LASV nsVGs produced in A549 cells. (a) Schematic of RNA-seq and bioinformatical analysis pipeline. (b-k) A549 cells were infected with LASV or TCRV (MOI = 0.1) for 48 hpi and total RNA was analysed by RNA-seq. (b) Relative continuous alignment rates to the LASV or TCRV reference genome compared to human alignments as determined by bowtie2. (c) Relative discontinuous alignment rates to the LASV or TCRV reference genome compared to total continuous viral alignments as determined by ViReMa. (d) Relative frequency of different nsVG types. (e) Relative frequency of origin and sense of LASV delVGs. (f) Relative frequency of origin and sense of TCRV delVGs. (g) Relative frequency of deletion sizes of LASV or TCRV delVGs. (h-i) Depiction and abundance of break and rejoin points of LASV delVGs stemming from the (h) L segment and (i) S segment. (j-k) Depiction and abundance of break and rejoin points of TCRV delVGs stemming from the (j) L segment and (k) S segment.

Analysis of junction-spanning reads in these two data sets revealed significantly higher relative levels in TCRV infected cells (Fig. 4c). Furthermore, LASV nsVGs exhibited a clear preference for genome duplications (dupVGs), while most TCRV nsVGs were generated through internal deletions (delVGs). Only a small fraction of nsVGs for both viruses exhibited copy-back (cbVGs) recombinations (Fig. 4d). Infections at high MOIs are known to often trigger aberrant genome replication and the synthesis of truncated viral genomes. We therefore infected A549 cells with LASV at a high MOI (MOI = 3) and measured viral replication and nsVG levels (Extended Data Figs. 3c-f). Total viral RNA levels increased between 4 and 24 hpi, but experienced a sudden decrease by 48 hpi (Extended Data Figs. 3c-d). This drop in total viral replication was not associated with an increase in either total nsVG (Extended Data Fig. 3e) or delVG levels (Extended Data Fig. 3f).

Since delVGs have been associated with the induction of antiviral host responses, we concentrated our following efforts on delVGs, specifically. While most LASV delVGs originated from the S segment (74%), the vast majority (95%) of TCRV delVGs stemmed from the L segment (Figs. 4e-f). There did not appear to be an obvious enrichment of delVGs in either the vRNA or cRNA sense for either virus. When analysing the size of the internal deletions across both segments, we observed that while there was a low frequency baseline of large deletions the majority of deletions were less than 40 nucleotides in size (Fig. 4g). In particular for TCRV there appeared a distinct high frequency peak in deletion size between 25 and 30 nucleotides in size, accounting for almost the entirety of delVG truncations. An investigation of the break and rejoin coordinates of these internal deletions and their respective abundances demonstrated clear patterns for LASV and TCRV (Figs. 4h-k): for all segments low abundance deletions of only a few nucleotides (*i.e.* the break and rejoin coordinates are very close together and therefore cluster as a diagonal line across the graph) could be observed across the entire genome segment, aligning with our findings that most delVGs are formed by small deletions (Fig. 4g). Furthermore, there is another accumulation of deletions that cluster towards the 5’ and 3’ ends of genome segments (*i.e.* the coordinates cluster in the top left or bottom right corners of the graph), representing delVGs that only consist of the genome termini and UTRs. In addition to these general observations, we also identified a few high frequency deletions in the LASV S segment (Fig. 4i) and the TCRV L segment (Fig. 4j). In the LASV S segment two clusters of abundant deletions were found: the first cluster of deletions was found in the exonuclease domain (exoN) of NP (nucleotides 2034 – 2234 of LASV S vRNA), whereas the second cluster was located in the IGR and the C-terminal region of NP (nucleotides 1461 – 1784 of LASV S vRNA). In the TCRV L segment a highly abundant cluster of deletions was found in the IGR (nucleotides 378 - 408 of the TCRV L vRNA). Prediction of the RNA secondary structure of the wild-type TCRV L segment IGR with or without the most abundant TCRV L segment deletion revealed a slightly changed IGR structure, which could potentially have an impact on viral mRNA transcription and gene expression (Extended Data Fig. 4a). To validate the presence of these small IGR deletions, A549 cells were infected with a separately generated TCRV stock and RT-PCR was performed around the TCRV L segment IGR to reveal a shorter PCR product in addition to the expected full-length product (Extended Data Fig. 4b). When performing virus- and segment-specific RT-PCR of entire virus segments we did not detect any major PCR products indicating truncated genomes (Extended Data Fig. 4c), which is in line with our bioinformatical analyses showing no major nsVGs with large deletion sizes (Figs. 4g-k). However, in line with previous reports we were able to observe several major truncated TCRV genome segments after five passages in Vero E6 cells (Extended Data Fig. 4d).

In summary, our data clearly indicates that unpassaged TCRV and LASV generate low levels of nsVGs already during primary infections which can accumulate over time and passages, but that there are distinct differences in the patterns and abundance of these produced nsVG for both viruses.

### Antiviral host response is mediated by RIG-I sensing and small aberrant viral RNAs

Previously, different pattern recognition receptors (PRRs) have been implicated in sensing of arenavirus infections, including RIG-I and PKR^14–16^ (Fig. 5a). Since we previously demonstrated that the induction of the antiviral host response requires active replication, we next investigated how RIG-I and PKR shape the innate immune response to OWA and NWA infections.

**Figure 5:**
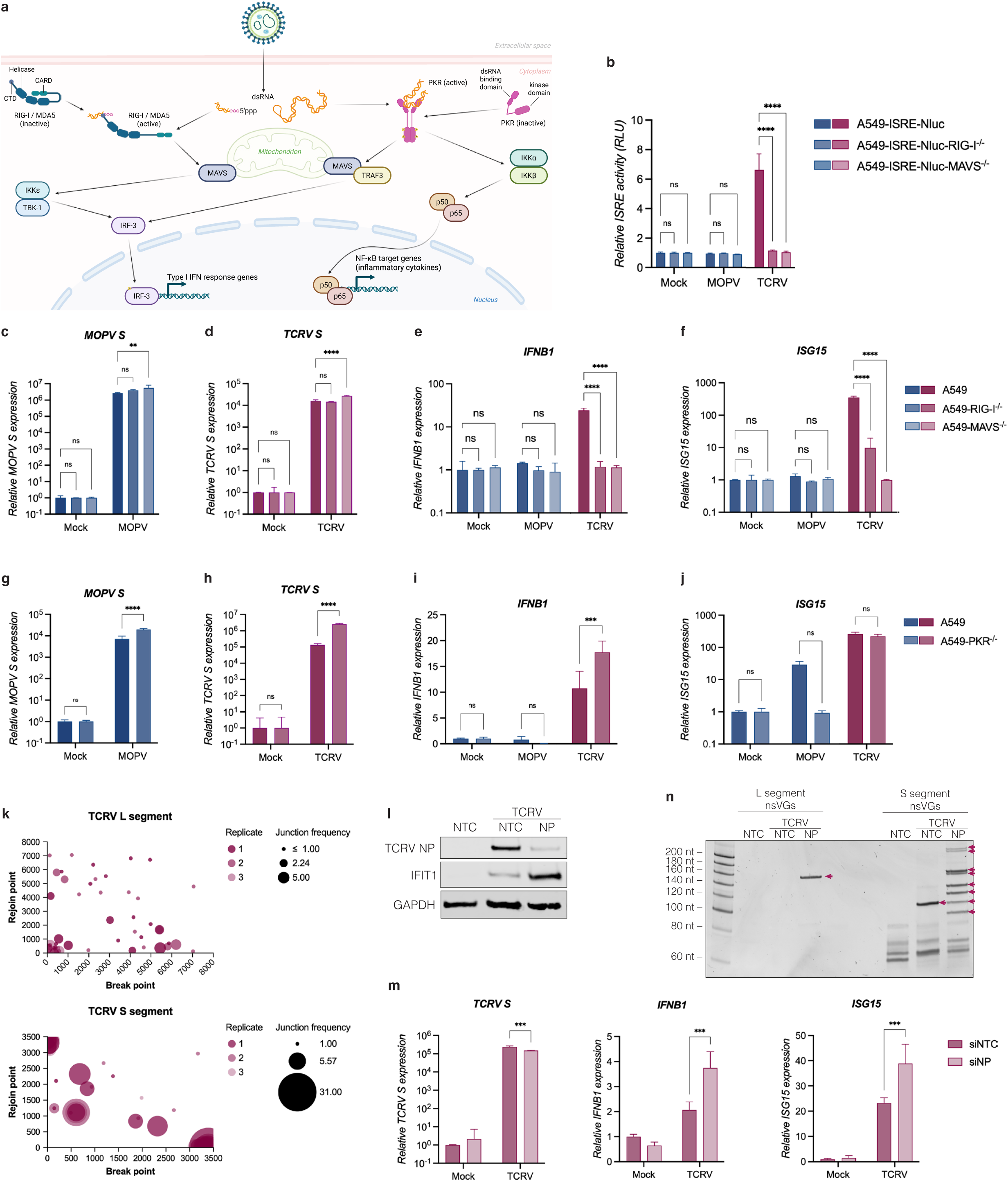
IFN response is mediated by RIG-I sensing of viral RNA and small aberrant replication products. (a) Schematic of viral RNA sensing and cellular antiviral signalling pathways. (b-f) A549-ISRE-Nluc2AGFP cells with or without RIG-I or MAVS knockouts were infected with TCRV or MOPV (MOI = 0.1). At 24 hpi ISRE-induced NanoLuc luciferase activity was measured or RNA was extracted and analysed by RT-qPCR. (b) The graph depicts the mean relative light units normalised to uninfected control cells from three independent biological replicates with error bars representing the standard deviation of the mean. (c) Extracted RNA was analysed by RT-qPCR for relative MOPV S RNA, (d) TCRV S RNA, (e) *IFNB1* and (f) *ISG15* gene expression. (g-j) A549-Cas9 cells with or without PKR knockouts were infected with TCRV or MOPV (MOI = 0.1) for 24 hpi. Extracted RNA was analysed by RT-qPCR for relative (g) MOPV S RNA, (h) TCRV S RNA, (i) *IFNB1* and (j) *ISG15* gene expression. (k) A549 cells were infected with LASV or TCRV (MOI = 0.1) for 48 hpi and extracted RNA was analysed by small RNA-seq. The graphs depict the coordinates and abundance of break and rejoin points of small TCRV delVGs. (l) A549 cells were transfected with siRNAs targeting the TCRV NP mRNA transcripts, followed by infection with TCRV (MOI = 0.1) for 24 hpi. Whole cells lysates were analysed by western blot for TCRV NP, IFIT1 and GAPDH protein expression. (m) Extracted RNA was analysed by RT-qPCR for the relative expression TCRV S RNA, *IFNB1* and *ISG15* gene expression. (n) The small RNA fraction (<200 nt) was analysed by vgRT-PCR and denaturing 7M urea 8% PAGE. Red arrows depict TCRV-derived mini viral RNAs. The graphs related to RT-qPCR data depict the mean fold change in gene expression normalised to the *ACTB* housekeeping gene from three independent biological replicates, with error bars representing the standard deviation of the mean. Statistical significance was determined by Two-way ANOVA with post-hoc Dunnett’s test (for b-f) and Šídák’s test (for g-j and m) for multiple comparisons: **** (*p* < 0.0001), *** (*p* < 0.001), ** (*p* < 0.01), * (*p* < 0.05), ns (*p* > 0.05).

First, we infected A549-ISRE-Nluc2AGFP cells with *RIG-I* or *MAVS* knockouts with MOPV or TCRV, respectively. At 24 hpi, we measured ISRE-mediated reporter activity and did not observe any significant activity in MOPV infected cells (Fig. 5b). In contrast, in wild-type cells infected with TCRV we could detect robust reporter activity, which was fully abrogated to control levels in the absence of *RIG-I* or *MAVS*. These protein-based reporter activity data were validated at the transcriptional level using RT-qPCR (Figs. 5c-f). Both MOPV (Fig. 5c) and TCRV (Fig. 5d) exhibited significantly increased levels of viral RNA replication in the absence of *MAVS*. Interestingly, knockout of *RIG-I* alone was not sufficient to increase viral replication. Transcriptional activity of the *IFNB1* (Fig. 5e) and *ISG15* (Fig. 5f) genes exhibited similar results as the reporter assay: while MOPV infection did not induce transcription of either gene, TCRV infection resulted in significant transcriptional activity, which was significantly or completely abrogated in the absence of *RIG-I* or *MAVS*.

Next, we infected A549-Cas9-PKR^-/-^ cells with MOPV or TCRV. Both MOPV (Fig. 5g) and TCRV (Fig. 5h) showed significantly higher levels of viral RNA after 24 hpi in the absence of *PKR*. In a similar manner to our previous data, TCRV, but not MOPV, induced robust levels of *IFNB1* transcription (Fig. 5i). Interestingly however, in the absence of *PKR*, transcriptional activity was significantly enhanced in TCRV infected cells. For *ISG15*, low levels of transcription could be observed in MOPV and high levels in TCRV infected cells and knockout of *PKR* had no significant effect on *ISG15* induction (Fig. 5j).

Given the clear indication that RIG-I is critical to viral RNA sensing and the fact that small viral RNAs have previously been shown to be key drivers of innate immune sensing of viruses^5^, we performed small RNA sequencing of TCRV infected A549 cells. Here, we were able to demonstrate that TCRV generates small aberrant viral RNAs that contain non-canonical junction sites preferentially in the S segment (Fig. 5k). While the coordinates of these junction sites are also found in more central regions of the segment, there appears to be a clear cluster of break and rejoin points at the extreme genome termini. This suggests that these reads are derived from highly truncated nsVG which consist only of the extreme terminal promoter and UTRs, which are highly complementary to each other and could form small dsRNA ligands for RIG-I and other PRRs. In order to validate these findings we used a previously established strategy to induce aberrant genome replication by negative-sense RNA viruses by restricting the availability of NP, which is an essential elongation factor for the viral replicase complex^17^. Using siRNA-mediated targeting of NP mRNA, we were able to effectively reduce viral NP expression (Fig. 5l), resulting in a small but significant reduction in viral RNA levels early in the infection cycle (Fig. 5m). Despite its reduced ability to replicate efficiently, knockdown of NP correlated with significant increases in IFIT1 protein expression (Fig. 5l), and *IFNB1* and *ISG15* transcriptional activation (Fig. 5m). This upregulation in bona fide antiviral genes strongly correlates with increased and more varied expression of small delVGs (Fig. 5n). These delVGs consist of both genome termini with large internal deletions, resulting in small aberrant viral RNAs of less than 200 nt in size.

Together, these data indicate that both RIG-I and PKR play important, yet distinct, roles in the sensing of arenavirus infections and the subsequent activation of the host antiviral response and in the control of viral replication. RIG-I appears to sense small aberrant viral genomes produced by TCRV that could form small blunt-ended dsRNA products.

## DISCUSSION

Arenaviruses pose a severe threat to global health and members of the virus family, including LASV, are listed as priority pathogens by the WHO R&D Blueprint for Epidemics^18^, by the Coalition for Epidemic Preparedness Innovations (CEPI)^19^ and the UK Health Security Agency (UKHSA)^20^. Currently, there are no licenced vaccines – apart from the live-attenuated Candid#1 against JUNV – and only the broad-spectrum antiviral Ribavirin, which has limited efficacy and significant side effects in humans, available^21–23^. It therefore remains important to gain a better understanding of the underlying molecular mechanisms that cause and exacerbate viral disease in humans.

Here, we demonstrate that there is a clear difference in the innate immune response of relevant human cell types to NWA and OWA. All NWA used in this study, comprising members from all four clades (A-D), uniformly induce a robust innate immune response, characterized by type I and II IFN, pro-inflammatory cytokines and activation of apoptosis pathways. In contrast, all tested OWA demonstrated a remarkable lack of antiviral response. Comprehensive transcriptional analysis of LASV infected cells by RNA sequencing only revealed a few significantly affected genes at any given time point, which is in stark contrast to our findings for TCRV, a prototypic NWA closely related to highly pathogenic NWA such as Junín virus (JUNV) and Machupo virus (MACV).

Previously, it was already demonstrated that NWA, including TCRV and the highly pathogenic JUNV and MACV, induce a strong innate immune response in human cells^15,24^. However, there have only been few studies detailing the effects of NWA on human cells in a comprehensive manner. Interestingly, highly pathogenic NWA have also been shown to be able to induce interferon-dependent pathogenic disease in mice and in non-human primates (NHP) characterized by cytokine storm-like features^25,26^. There is only limited data about apathogenic NWA in NHP, but infection with TCRV does not induce any signs of viral disease in marmosets^27^. A limitation of our study is that we were not able to include any highly pathogenic NWA: however, given the previously published findings that JUNV and its close relative TCRV exhibit similar levels of innate immune induction in cell culture, it is reasonable to assume that highly pathogenic New World viruses would have exhibited a similar phenotype in our study^15^.

This contrasts with previous reports that infection of primary human^24,28,29^ and Natal multimammate mice (the natural rodent host species for LASV and MOPV)^30–32^ cells with LASV, and to a lesser degree MOPV and MOBV, failed to induce a significant antiviral host response, confirming our own data presented in Figures 1 and 2. However, in NHP models LASV exhibits strain- and lineage-dependent clinical phenotypes: fatal LASV infections are associated with systemic viral replication, a defective and dysregulated immune response, cytokine storms and multi-organ failure, whereas non-fatal LASV infections appear to be effectively controlled locally restricted by an appropriate immune response^33–37^. Similar to non-fatal LASV infections, challenge with the apathogenic MOPV appears to be rapidly and effectively controlled in NHP and does not lead to pathogenic disease^38^. Together, these data suggest that on the one hand there exists a fundamental difference in NWA and OWA innate immune induction in a relatively simple and homogenous biological system such as cell culture. On the other hand, in complex NHP models there appears a clear distinction in disease manifestation between highly pathogenic and apathogenic viruses. How these divergent phenotypes can fully be reconciled remains yet to be determined.

In our study we clearly demonstrate that RIG-I and MAVS together play a fundamental and essential role in arenavirus sensing and restriction. This is in line with previous reports that highlight the critical role of RIG-I in the antiviral response to arenaviruses^14,39–41^. In addition, we also show that PKR plays an important role in regulating the broader antiviral response to arenaviruses but is dispensable for the induction of the initial type I/III IFN response.

Previously, it had been suggested that there are fundamental differences in the abilities of arenavirus proteins to antagonise the IFN signalling pathway^40,41^. In our study we were unable to detect any significant differences in the IFN antagonistic potential of both NP and Z proteins of a wide range of arenaviruses. In addition, in the context of virus infection we also showed that both LATV and MOPV (to a lesser extent) were able to inhibit a poly(I:C) induced IFN response, but not a type I IFN induced host response, also suggesting that there is no significant difference in the ability of NWA and OWA to suppress the activation of the antiviral host response in a manner consistent with the distinct infection phenotypes. Analysis of the relative gene expression levels of the individual ORFs of LASV and TCRV revealed some differences in relative NP and Z expression levels between both viruses, with LASV exhibiting higher NP and TCRV higher Z expression levels at 48 hpi. However, further studies are required to address whether viral IFN antagonist expression levels are generally different between different groups of arenaviruses.

The accumulation of nsVGs has been associated with increased disease severity for multiple viruses, including IAV^5^, SARS-CoV-2^9^ and RSV^8^. In a recent study in African green monkey cells using Illumina-based short-read sequencing, it was demonstrated that arenaviruses, including LCMV, JUNV and Parana virus, can generate delVGs^12^. These internal truncations tend to either omit large parts of entire genome segments or ORFs, or alternatively cluster in and around the highly structured IGR sequences between the ORFs. These IGR deletions were shown to be virus-specific and independent of any biases potentially introduced by reverse transcription or PCR. In line with these data, we observe a very similar pattern where we see clusters of nsVG junctions either close to the genome termini or around the IGR sequences. While our data supports delVG formation for TCRV even in unpassaged virus stocks, the vast majority of these internal deletions are very small in size (Fig. 4h) and are concentrated around the L segment IGR (Fig. 4k). However, after only five passages in Vero E6 cells we were able to see highly abundant significantly truncated genomes for TCRV, which was also observed in another recent study of serially passaged TCRV using amplicon-based Nanopore sequencing, where the authors not only were able to identify similar truncation patterns using different experimental and analysis methods, but also more complex aberrant viral RNAs with multiple breakpoints and duplications of promoter regions^13^. Interestingly, we were also able to identify a large proportion of genome duplications across the entire genome in particular for LASV (and to a much lesser degree for TCRV), which were not detected previously due to limitations of the used *VODKA2* bioinformatical pipeline to detect genome duplications^42^. To address the mechanistic role and consequence of these specific truncations and duplications more work is required, but it is likely that these recombinations can have a profound impact on viral gene expression, as demonstrated for some IGR truncations observed in LCMV infections^12^.

Given the critical role of RIG-I/MAVS in sensing and controlling both OWA and NWA infections (Fig. 5b-f), our findings that TCRV generates small (<200 nt) aberrant replication products is highly significant. Analysis of break and rejoin points of these aberrant replication products highlighted abundant genome truncations that result in mini viral RNAs (mvRNA) consisting of both extreme genome termini only (Figs. 5k and 5n). These resulting mvRNAs are highly reminiscent of mvRNAs identified in IAV infections and which have been shown to be correlated to the induction of the innate immune response and the cytokine storm associated with highly pathogenic avian influenza virus strains^5–7,17^. Due to the extreme sequence complementarity of the 5’ and 3’ genome termini of arenaviruses, these mvRNAs could potentially fold up on themselves and form small blunt-ended dsRNA products that might serve as high-affinity ligands for RIG-I. Alternative mechanisms for RIG-I activation observed for IAV involve mvRNA hybridization with complementary transcription or replication products of these mvRNAs, resulting in longer dsRNA complexes that can serve as ligands for RIG-I detection^7^. It is important to note that arenaviruses are known to generate 5’ triphosphate ends, which are only very inefficiently captured using the library preparation method for small RNA sequencing used in this study. A ligation-independent approach to small RNA sequencing will likely result in much more efficient capture of mini arenavirus RNAs. In our previous publication we detailed an efficient strategy to coax negative-sense RNA viruses into preferentially generating small aberrant replication products over full-length genome equivalents using an siRNA-mediated NP depletion approach^17^. Using the same strategy, we were able to demonstrate that an increase in variety and abundance of TCRV mvRNAs is highly correlated to an increased antiviral host response (Figs. 5l-m). Due to the fact that arenavirus NP is not only an essential co-factor of the viral replication machinery, but also a critical IFN antagonist, it is difficult to separate the effects of mvRNA generation and the loss of NP-mediated IFN antagonism^43,44^. Nevertheless, it remains obvious, that the reduced availability of TCRV NP results in less full viral genome replication, but more small aberrant and potentially dsRNA products (whether that is due to increased mvRNA generation or reduced dsRNA exonuclease activity by NP) which is responsible for a significantly increased host cellular response. It remains to be seen whether synthesis of aberrant mvRNA products might help to explain differences in innate immune induction and pathogenesis by NWA and OWA. But considering the proven presence of these mvRNA-like ligands in NWA infections, it is tempting to speculate that these aberrant replication products could be – as in highly pathogenic avian influenza - key drivers of the cytokine storm and highly inflammatory viral disease observed in animal models and patients infected with highly pathogenic arenaviruses^26^.

Overall, we identified distinct patterns in innate immune induction by NWA and OWA, respectively, and demonstrated that these phenotypes can be attributed to viral replication products and not to significant differences in the ability of viruses to inhibit RNA sensing or IFN signalling. The activation of antiviral responses was clearly initiated by RIG-I sensing and aided by PKR. In addition, we characterized defective viral genome replication of representative OWA and NWA, highlighting differences in nature and abundance of nsVG production and identified aberrant mvRNA species that are correlated to the induction of the IFN response.

## MATERIALS AND METHODS

### Cell culture

Human alveolar basal epithelial carcinoma cells (A549; ATCC, CCL-185), human embryonic kidney cells (HEK-293T; ATCC, CRL-3216), African green monkey kidney epithelial cells (Vero E6; ATCC, CRL-1586) and Syrian golden hamster kidney fibroblast cells (BHK-21; ATCC, CCL-10) were commercially obtained. Human pulmonary microvascular endothelial cells (HPMEC; PromoCell) were commercially obtained and maintained in complete Endothelial Cell Growth Medium MV (PromoCell) and used at low passage numbers (<10). A549-ISRE-mScarlet cells encoding an mScarlet fluorescent reporter under the control of an interferon-stimulated response element (ISRE) were a kind gift by Charles Rice (Rockefeller University). A549-Cas9 and A549-Cas9-PKR^-/-^ cells have previously been described and were a kind gift by Linda Brunotte (University of Münster)^45^. A549-ISRE-Nluc2AGFP, A549-ISRE-Nluc2AGFP-DDX58^-/-^ and A549-ISRE-Nluc2AGFP-MAVS^-/-^ cells encoding a NanoLuc luciferase and GFP reporter under the control of an interferon-stimulated response element (ISRE) as well as genetic knockouts for *DDX58* or *MAVS* were a kind gift by Andrew Mehle (University of Wisconsin-Madison) and have previously been described^46^. Unless stated otherwise, all cells were maintained in Dulbecco’s Modified Eagle Medium (DMEM) supplemented with 10% FBS, L-Glutamine, MEM Non-Essential Amino Acids, 100 U/ml penicillin and 100 μg/ml streptomycin and cultured at 37°C and 5% CO_2_.

### Virus culture and infections

Tacaribe virus (TCRV; *Mammarenavirus tacaribeense*) strain TRVL-11573 was obtained from ATCC (VR-1556). Pichinde virus (PICV; *Mammarenavirus caliense*) strain CoAn-3739, Latino virus (LATV; *Mammarenavirus latinum*) strain MARU-10924 and Tamiami virus (TAMV; *Mammarenavirus tamiamiense*) strain W-10777 were obtained through the NIH Biodefense and Emerging Infections Research Resources Repository, NIAID, NIH. Lymphocytic choriomeningitis virus (LCMV; *Mammarenavirus choriomeningitidis*) strain Armstrong 53b was a kind gift by Anke Kraft (Twincore Centre for Experimental and Clinical Infection Research, Hannover, Germany). Lassa virus (LASV; *Mammarenavirus lassaense*) strain Bantou 366 (Ba366) was a kind gift by the Institute of Virology at the University of Marburg^47^. Mopeia virus (MOPV; *Mammarenavirus mopeiaense*) strain AN-21366-BNI was originally obtained from National Collection of Pathogenic Viruses, Porton Down, Salisbury, UK^48^. Mobala virus (MOBV; *Mammarenavirus praomyidis*) strain 3099 was originally obtained from the Centers for Disease Control and Prevention, Atlanta, USA^49^.

LASV was propagated and titrated by immunofocus assay on BHK-21 cells as previously described^50^. All other viruses were propagated in Vero E6 cells in DMEM supplemented with 2% FBS, L-Glutamine, MEM Non-Essential Amino Acids, 100 U/ml penicillin and 100 μg/ml streptomycin. Virus-containing cell culture supernatants were collected 4 to 7 days post-infection (dpi) and purified from small soluble cell culture contaminants and concentrated using Amicon Ultra-15 centrifugal filter units (100-kDa molecular weight cut-off). Infectious virus titres were determined by immuno-plaque assays. Briefly, Vero E6 cells were infected with sequential 10-fold dilutions of virus and overlayed with 1.2% Avicel CL-611 in MEM containing 2% FBS, 10 mM HEPES, MEM Non-Essential Amino Acids and 100 U/ml penicillin and 100 μg/ml streptomycin. At 7 days post-infection, cells were fixed in 3.7% formaldehyde and permeabilized in 0.5% Triton X-100 and immunostained with the primary antibodies detailed in Table 1. Infected cells were visualized with an IRDye-conjugated anti-mouse IgG (IRDye 800CW, 926-32210) secondary antibody. Fluorescent signal of stained plaques was detected using an Odyssey CLx imaging system (LI-COR) and analysed with the Image Studio software (LI-COR).

**Table 1.** Antibodies used for immuno-plaque assays.

| **Antibody** | **Clone** | **Species** | **Source / Reference** |
| --- | --- | --- | --- |
| *LASV NP mAb* | L2B5 | MOBV, MOPV | Hufert *et al.* (1989) <sup>51</sup> |
| *LASV NP mAb* | L2F1 | LASV | Hufert *et al.* (1989) <sup>51</sup> |
| *LCMV mAb* | M104 | LCMV | Progen (651106) |
| *JUNV NP mAb* | IC06-BE10 | TCRV, TAMV | BEI Resources (NR-43777) |
| *JUNV NP mAb* | JB02-BF08 | LATV, PICV | BEI Resources (NR-40328) |

All LASV infection were performed in DMEM supplemented with 5% FBS, and at the time of harvest cells were washed once in PBS, dissociated using Trypsin-EDTA solution and inactivated as appropriate for the required analyses. All other virus infections were performed in DMEM supplemented with 2% FBS, L-Glutamine, MEM Non-Essential Amino Acids, 100 U/ml penicillin and 100 μg/ml streptomycin.

All work involving infectious LASV was performed at the BSL-4 facility of the Bernhard Nocht Institute for Tropical Medicine (Hamburg, Germany) in accordance with their institutional biosafety requirements.

### Serial virus passaging

In order to serially passage TCRV, Vero E6 were infected with TCRV at an MOI of 1 in infection media. Every 3-4 days, virus-containing supernatant was transferred onto naïve Vero E6 cells in infection media for a total of five blind passages. After passage 5 (p5) RNA was extracted from the infected Vero E6 cells and analysed by vgRT-PCR as detailed below.

### siRNA-mediated silencing of NP expression

Custom siRNAs were designed against the NP mRNA sequence of TCRV and ordered from Integrated DNA Technologies. Approximately 240,000 A549 cells were transfected with 10 pmol of nontargeting control siRNA or NP targeting siRNA using Lipofectamine RNAiMax and Opti-MEM according to the manufacturer’s instructions. 24 h post-transfection, cells were infected with TCRV at an MOI of 0.1 in complete DMEM supplemented with 2% FBS. At 24 hpi, whole cell lysates and total RNA was extracted and processed as described below.

### Western blot

For western blot analysis, whole-cell lysates were obtained through cell lysis in RIPA buffer supplemented with 1% SDS and sonication. Lysates were analysed by SDS-PAGE and transferred onto nitrocellulose membranes. Target proteins were detected using primary antibodies again Actin (Invitrogen; MA5-11869), GAPDH (Sigma-Aldrich; G9545), IFIT1 (Cell Signaling Technology; 14769), STAT1 (Cell Signaling Technology; 9172), p-STAT1 [Tyr701] (Cell Signaling Technology; 9167), MX1 (Abcam; ab95926), LASV NP (GeneTex; GTX636832), and JUNV NP (BEI Resources; NR-48834). Primary antibodies were detected using IRDye-conjugated anti-mouse IgG (IRDye 680RD, 926-68070; IRDye 800CW, 926-32210) and anti-rabbit IgG (IRDye 680RD, 926-68071; IRDye 800CW, 926-32211) secondary antibodies. Fluorescent signals were detected using an Odyssey CLx imaging system (LI-COR) and analysed with the Image Studio software (LI-COR).

### Quantitative real-time PCR

Cells were lysed in TRIzol Reagent (Invitrogen) and total RNA was extracted and DNase I-treated using the Direct-zol RNA Miniprep kit (Zymo Research) according to the manufacturer’s instructions. Reverse transcription using oligo(dT)/random hexamer primers was performed using the PrimeScript RT Master Mix (TaKaRa Bio) according to the manufacturer’s instructions. Quantitative real-time PCR was performed using the TB Green Premix Ex Taq II (Tli RNase H Plus) master mix (TaKaRa Bio) or the Luna Universal qPCR Master Mix (NEB) on a LightCycler 480 Instrument (Roche) according to the manufacturers’ instructions. Primer sequences used are detailed in Table 2. Delta-delta-cycle threshold (ΔΔ*C_T_*) values were determined relative to a housekeeping gene (GAPDH or ACTB) and normalized to the mean of the indicated control group.

**Table 2.**
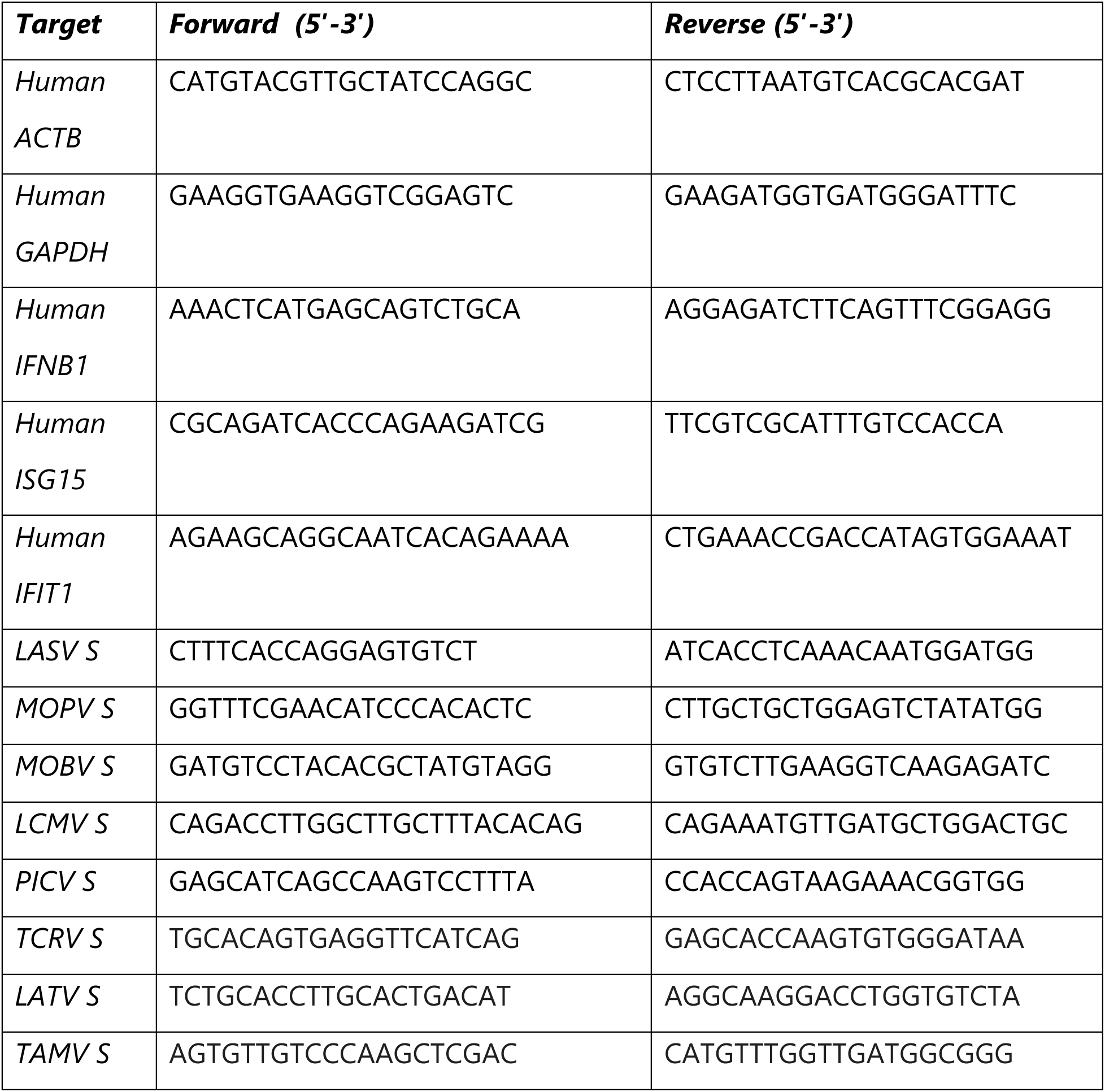
Primers used for qPCR and RT-PCR.

### Immunofluorescence assay

For MOBV infections approximately 10,000 A549 cells were infected with MOBV at an MOI of 0.1 in DMEM supplemented with 2% FBS, L-Glutamine, MEM Non-Essential Amino Acids, 100 U/ml penicillin and 100 μg/ml streptomycin. At 48 hpi, cells were washed once in PBS and fixed in 4% formaldehyde. For LASV infections approximately 5,000 A549 cells were infected with LASV at an MOI of 0.1 in DMEM supplemented with 5% FBS. At 48 hpi, cells were washed once in PBS, dissociated with Trypsin-EDTA solution, mounted on glass cover slides and fixed in chilled 100% acetone. Fixed cells were permeabilised in 0.5% Triton X-100 and blocked in 5% BSA dissolved in PBS. Viral NP was detected using a monoclonal cross-reactive LASV NP antibody (clone L2B5) and an Alexa Fluor Plus 488 Highly Cross-Adsorbed Goat anti-Mouse IgG secondary antibody (Invitrogen, A32723) or an Alexa Fluor 568 Highly Cross-Adsorbed Donkey anti-Mouse IgG secondary antibody (Invitrogen, A10042). Cell nuclei were counterstained with 4′,6-diamidino-2-phenylindole (DAPI). Images were captured on an EVOS M5000 Imaging System (Invitrogen).

### Live-cell imaging

Approximately 4,000 A549-ISRE-mScarlet cells per well were seeded in 96-well plates. On the following day, cell culture media was removed, and cells were infected with the indicated viruses at an MOI of 0.1 in DMEM without phenol red supplemented with 2% FBS. Live-cell imaging was performed using an IncuCyte SX5 Live-Cell Analysis System (Sartorius) using the 10X objective and orange optical module (400 ms) and the adherent cell-by-cell imaging settings. Four images per well were acquired at 4 h intervals for 5 dpi. Following live-cell imaging, background subtraction and image analysis was performed using the IncuCyte 2024A GUI software (Sartorius) to calculate mean fluorescence intensity per image.

### UV inactivation of TCRV

For UV-inactivation of TCRV, 0.5 ml of virus stock was placed in a sterile cell culture dish and UV cross-linking (254 nm, 1 J/cm^2^) of the virus stock was performed using a CX-2000 UV Crosslinker (UVP). Subsequently, A549 cells were infected for 72 h at an MOI of 0.01 with infectious untreated TCRV or equivalent volumes of UV-inactivated virus. At the time of samples collection, cells were lysed and collected for western blot analysis as described above.

### Plasmids

Plasmids pcDNA3a^52^ and pGL3-IFNB1^53^ have previously been described. Plasmids pCAGGS-(FLAG)RIG-I and pCAGGS-(FLAG)mScarlet were generated by In-Fusion cloning of the mScarlet or RIG-I coding sequences into an empty pCAGGS vector with an N-terminal FLAG-tag using the In-Fusion Snap Assembly Master Mix (TaKaRa Bio). Plasmids pcDNA-LASV-NP, pcDNA-MOBV-NP, pcDNA-LCMV-NP, pcDNA-TCRV-NP, pcDNA-PICV-NP, pcDNA-TAMV-NP, pcDNA-LASV-Z, pcDNA-MOBV-Z, pcDNA-LCMV-Z, pcDNA-TCRV-Z, pcDNA-PICV-Z and pcDNA-LATV-Z were generated by In-Fusion cloning. Here, RNA was extracted from infected cells using TRIzol Reagent (Invitrogen) and the Direct-zol RNA Miniprep kit (Zymo Research) according to the manufacturer’s instructions. RNA was reverse transcribed into cDNA using the PrimeScript RT Master Mix (TaKaRa Bio) and the respective NP and Z open reading frames for each virus were subsequently amplified using the CloneAmp HiFi PCR Premix (TaKaRa Bio). In-Fusion assembly into an empty pcDNA3a vector was performed using the In-Fusion Snap Assembly Master Mix (TaKaRa Bio) according to the manufacturer’s instructions. Plasmid pcDNA-IAV-NS1 was generated by In-Fusion cloning. Here, the NS1 open reading frame was amplified from the pHW2000-IAV-WSN-NS plasmid^54^ using the CloneAmp HiFi PCR Premix (TaKaRa Bio). In-Fusion assembly into an empty pcDNA3a vector was performed using the In-Fusion Snap Assembly Master Mix (TaKaRa Bio) according to the manufacturer’s instructions. Plasmid pcDNA-(FLAG)mScarlet was generated by In-Fusion cloning. Here, a codon-optimised N-terminally FLAG-tagged mScarlet gene was synthesised (Integrated DNA Technologies) and In-Fusion cloning into an empty pcDNA3a vector was performed using the In-Fusion Snap Assembly Master Mix (TaKaRa Bio) according to the manufacturer’s instructions. All plasmid sequences were verified by Sanger sequencing.

### Interferon reporter assays

For infection-based interferon reporter assays, approximately 40,000 A549-ISRE-Nluc2AGFP, A549-ISRE-Nluc2AGFP-RIG-I^-/-^ or A549-ISRE-Nluc2AGFP-MAVS^-/-^ cells per well in a 96-well plate were infected with TCRV or MOPV in complete DMEM supplemented with 2% FBS at an MOI of 0.1 for 24 hpi, as indicated. These cells express a NanoLuc luciferase and GFP reporter under the control of an interferon-stimulated response element (ISRE)^46^. For IFN inhibition assays, approximately 240,000 A549-ISRE-Nluc2AGFP cells per well in a 24-well plate were infected with LATV or MOPV in complete DMEM supplemented with 2% FBS at an MOI of 0.1 for 72 hpi before being treated with DMSO, recombinant 1,000 iu/ml IFN-ɑ2b, or 1 µg poly(I:C) RNA for 24 h. For RNA transfection-based interferon reporter assays, approximately 40,000 A549-ISRE-Nluc2AGFP cells in a 96-well plate were transfected for 24 h with 300 ng RNA or poly(I:C) using Lipofectamine RNAiMax and Opti-MEM in complete DMEM supplemented with 10% FBS. For plasmid transfection-based interferon reporter assays, approximately 40,000 HEK-293T cells in a 96-well plate were transfected for 24 h with 50 ng pGL3-IFNB1, 50 ng pCAGGS-(FLAG)RIG-I or pCAGGS-(FLAG)mScarlet, and 200 ng pcDNA-(FLAG)mScarlet, pcDNA-IAV-NS1, or pcDNA vectors expressing arenavirus NP or Z protein using Lipofectamine 2000 and Opti-MEM in complete DMEM supplemented with 10% FBS.

At the indicated time points, cell culture supernatants were removed, and cells were lysed in luciferase lysis buffer (1% Triton X-100, 25 mM Gly-Gly, 15 mM MgSO4, 4 mM EGTA, 1 mM DTT). Firefly and NanoLuc luciferase activities were measured using an inhouse protocol on a Centro Microplate Luminometer (Berthold Technologies).

### Viral genome RT-PCR (vgRT-PCR)

Total RNA was fractionated into small (<200 nt) and large (>200 nt) RNA fractions using the RNA Clean & Concentrator-5 kit (Zymo Research). Reverse transcription of the >200 nt RNA fraction was performed using the Maxima H Minus Reverse Transcriptase (Thermo Scientific) and primers complementary to the conserved terminal 5’ and 3’ promoter regions of all arenavirus L and S segments (Table 3). cDNA was subsequently purified from excess RT primers using the NucleoSpin Gel and PCR Clean-Up Kit (Macherey-Nagel) with a 20% NTI buffer according to the manufacturer’s instructions. Reverse transcription of the <200 nt RNA fraction was performed using the PrimeScript RT Master Mix (TaKaRa Bio) and oligo(dT)/random hexamer primers.

**Table 3.**
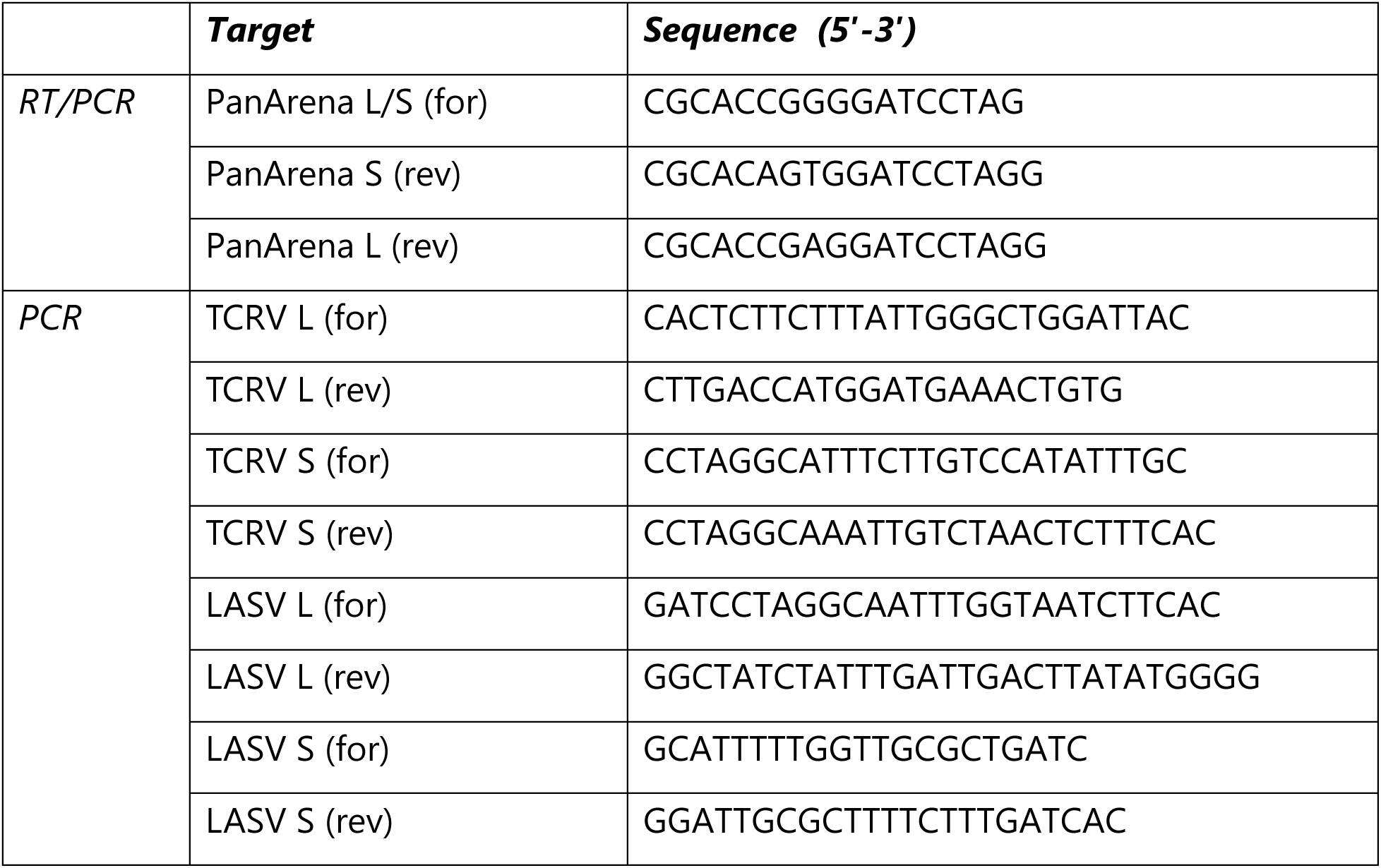
Primers used for vgRT-PCR.

PCR amplification of viral RNA segments with terminal promoter regions was performed using the PrimeSTAR GXL DNA Polymerase (TaKaRa Bio) and primers complementary to the 5’ and 3’ UTRs of the L and S segments (for >200 nt RNA) or primers complementary to the extreme termini used during RT (for <200 nt RNA) as detailed in Table 3. PCR reactions of >200 nt RNA fractions were resolved by 0.7% agarose gel electrophoresis. PCR reactions of <200 nt RNA fractions were resolved by 8 M urea 10% PAGE and stained using ethidium bromide. Gels were imaged on a GelDoc Go Gel Imaging System (Bio-Rad).

### RT-PCR

For the RT-PCR validation of IGR deletions, reverse transcription of viral RNA was performed as described above for vgRT-PCR. PCR analysis was performed using the PrimeSTAR GXL DNA Polymerase (TaKaRa Bio) and custom primers (for: 5’-CCATAAGAGTCCCGCTAGAG-3’, rev: 5’-CACAGTGGAGGACATTGAGTAG-3’). PCR products were analysed by denaturing 7 M urea 12% PAGE in TBE buffer and DNA bands were stained using ethidium bromide and imaged on a GelDoc Go Gel Imaging System (Bio-Rad).

#### RNA secondary structure prediction

RNA secondary structure prediction was performed using the UNAFold Web Server (https://www.unafold.org/) with default parameters^55^.

### Phylogenetic analysis of arenavirus RdRp sequences

The indicated nucleic acid sequences for the indicated arenaviruses were downloaded from GenBank and NCBI and the open reading frames for the viral RdRp proteins were translated using ExPASy (Swiss Institute of Bioinformatics)^56^. Phylogenetic analysis of arenavirus RdRp amino acid sequences was inferred using Maximum-Likelihood method and Jones-Taylor-Thornton model of amino acid substitutions and the tree with highest log likelihood is shown^57^. The initial tree for the heuristic search was selected by choosing the tree with the superior log-likelihood between a Neighbor-Joining (NJ) tree^58^ and a Maximum Parsimony (MP) tree. The NJ tree was generated using a matrix of pairwise distances computed using the Jones-Taylor-Thornton (1992) model^57^. The MP tree had the shortest length among 10 MP tree searches, each performed with a randomly generated starting tree. The analytical procedure encompassed 32 amino acid sequences with 2,338 positions in the final dataset. Evolutionary analyses were conducted in the Molecular Evolutionary Genetics Analysis (MEGA) software (version 12.1)^59,60^.

### RNA sequencing

Cells were lysed in TRIzol Reagent (Invitrogen) and total RNA was extracted and DNase I-treated using the Direct-zol RNA Miniprep kit (Zymo Research) according to the manufacturer’s instructions. Quality and integrity of total RNA was controlled on a 5200 Fragment Analyzer System (Agilent Technologies).

For total RNA sequencing, sequencing libraries were generated from 500 ng total RNA using the Ribo-off rRNA Depletion Kit (human, mouse, rat) (Vazyme BioTech Co.Ltd.) for rRNA depletion followed by the NEBNext Ultra II Directional RNA Library Prep Kit (New England BioLabs) according to manufactureŕs instructions. The libraries were treated with Illumina Free Adapter Blocking Reagent and were sequenced on an Illumina NovaSeq 6000 system using the NovaSeq 6000 S1 Reagent Kit (100 cycles, paired end run 2x 50 bp, 300 cycles, paired end run 2x 150 bp) with an average of 5 x10^7^ reads per RNA sample. For small RNA sequencing, sequencing libraries were generated from 500 ng total RNA using the Ribo-off rRNA Depletion Kit (human, mouse, rat) (Vazyme BioTech Co.Ltd.) for rRNA depletion followed by the NEBNext Multiplex Small RNA Library Prep Kit for Illumina (New England BioLabs) according to manufactureŕs instructions. The libraries were treated with Illumina Free Adapter Blocking Reagent and were sequenced on an Illumina NovaSeq X system using the NovaSeq X 1.5B Reagent Kit (300 cycles, paired end run 2x 150 bp) with an average of 1 x10^6^ reads per RNA sample. All library preparations and Illumina sequencing were conducted at the Genome Analytics (GMAK) core facility at the Helmholtz Centre for Infection Research.

### Bioinformatical analyses

Adapter and low complexity sequences were trimmed from raw sequencing fastq reads using the *fastp* software package (*v1.0.1*)^61^ and quality control was performed using the *fastqc* package (*v0.12.1*)^62^. Trimmed reads were aligned to the human transcriptome (GRCh38.p14, GENCODE release 49) using the *salmon* software (*v1.10.3*) in quasi-mapping mode^63^.

All statistical analyses of differentially expressed genes (DEGs) were performed in R using Bioconductor packages. Annotation and summarisation of transcripts to genes were carried out in R the *tximport* and *txdbmaker* packages^64^. Differential expression of genes (|log_2_FC| > 2 and adjusted *p-*value < 0.05) was performed using *DESeq2* (*v1.50.2*) with Benjamini-Hochberg (BH) correction for multiple testing^65^ using *ashr* adaptive shrinkage^66^. To diminish statistical noise, samples were always compared to relevant matching uninfected controls that were collected at the same time. Principal component analyses (PCA) and distance matrices were generated using the *DES*eq2 (*v1.50.2*) and *pheatmap* (*v1.0.13*) packages. Volcano plots were generated using the *EnhancedVolcano* package (*v1.28.2*). Gene set enrichment analysis (GSEA) was performed using the *clusterProfiler* (*v4.18.3*) and *enrichplot* (*v1.30.4*) packages. Heatmaps were constructed using the differentially expressed genes belonging to the Hallmark gene sets *Interferon Alpha Response* (M5911), *Interferon Gamma Response* (M5913), *Inflammatory Response* (M5932), and *Apoptosis* (M5902) using the *pheatmap* package (*v1.0.13*)^67^.

For alignments to viral genomes, trimmed fastq reads were aligned to the human genome (GRCh38.p14, GENCODE release 49) using *bowtie2* (*v2.5.4*) with default parameters^68^. Unaligned reads were subsequently aligned to the LASV strain Ba366 (GenBank accession numbers GU830839 and GU979513) or TCRV strain TRVL-11573 (GenBank accession numbers MT081316 and MT081317) genomes using *bowtie2* (*v2.5.4*) with default parameters^68^. Quantification of viral reads was performed using *samtools* (*v1.23*) and normalised to the number of human aligned reads. Virus genome coverage per nucleotide was determined using *bedtools* (*v2.31.1*). For identification of non-canonical junction sites and recombinations, reads that did not map to the human genome were analysed using the Virus Recombination Mapper (*ViReMa*) algorithm (*v0.29*) and the *bowtie* (*v1.3.1*) aligner with parameters *--Defuzz 0 --MicroInDel_Length 5 --Seed 15 --ErrorDensity 1,25*^69,70^. The resulting BED and BEDPE files were used to quantify and analyse recombination and deletion events.

### Statistical analyses

All other statistical analyses were performed as indicated in figure legends using GraphPad Prism version 10.6.1 for macOS, GraphPad Software, Boston, Massachusetts USA, www.graphpad.com.

## Data availability

The raw sequencing data sets generated in this study are available on the NCBI Sequence Read Archive (SRA) server with the BioProject ID xxx.

## ACKNOWLEDGEMENTS

A.A. was supported by the Hannover Biomedical Research School (HBRS) and the Center for Infection Biology (ZIB). This study was also funded by the Deutsche Forschungsgemeinschaft (DFG, German Research Foundation) under Germany’s Excellence Strategy - EXC 2155 - project number 390874280 (to B.E.N. and T.P.) and the Swedish Research Council (2023-02595 and 2024-03783 to B.E.N.).

The following reagent was obtained through BEI Resources, NIAID, NIH: Monoclonal Anti-Junin Virus, Clone NA05-AG12 (produced *in vitro*), NR-48834. Monoclonal Anti-Junin Virus, Clone JB02-BF08 (produced *in vitro*), NR-40328. Monoclonal Anti-Junin Virus, Clone IC06-BE10 (produced *in vitro*), NR-43777. Tamiami Virus, W-10777, NR-10178. Latino Virus, MARU 10924, NR-12236. Pichinde Virus, CoAn-3739, NR-10177.

---

**Extended Data Figure 1:**
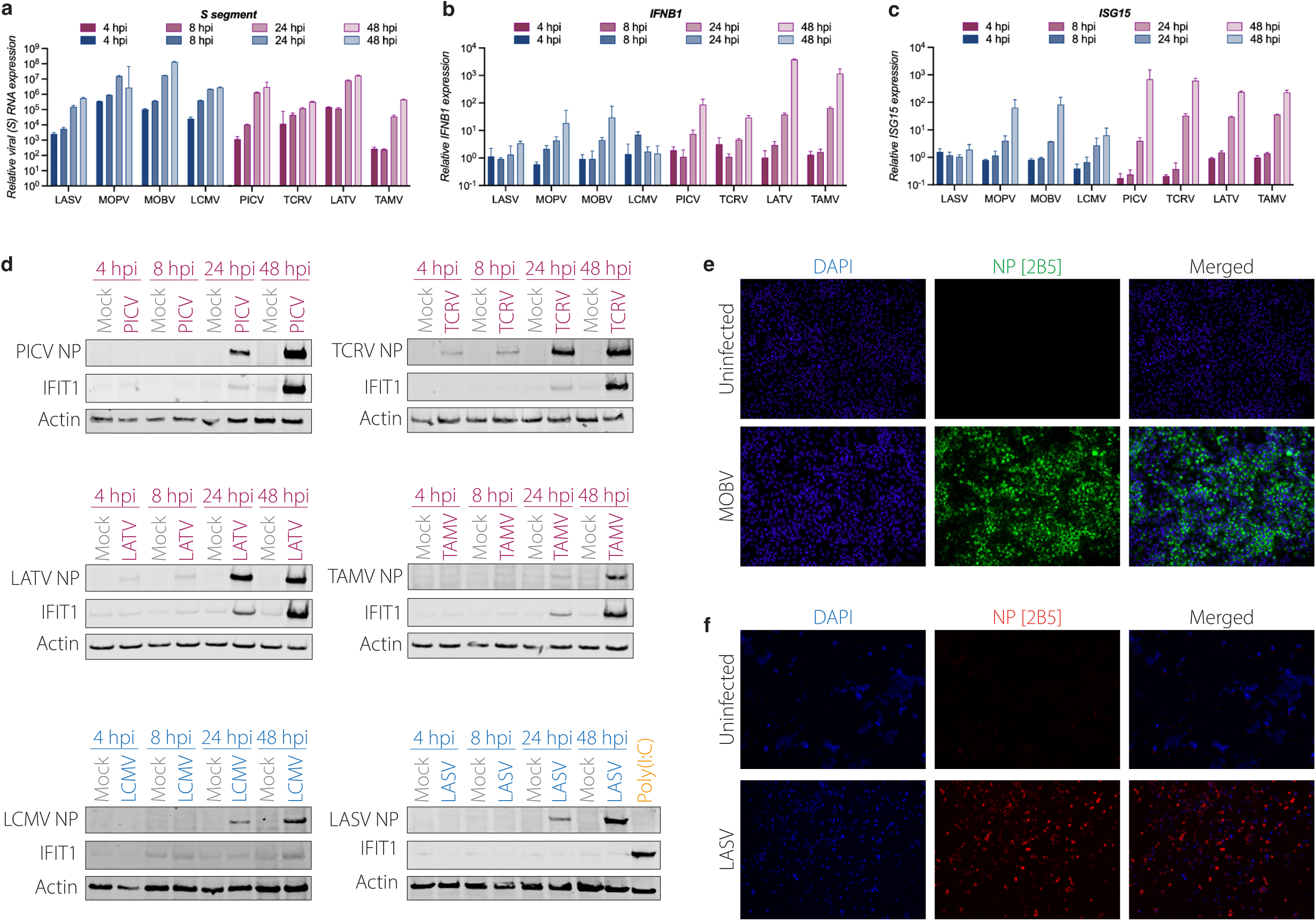
NWA induce a robust type I IFN response. (a-d) A549 cells were infected with the indicated arenaviruses at an MOI of 0.1. At the indicated times post infection RNA was extracted and analysed by RT-qPCR for the relative gene expression of (a) viral S segment RNA, (b) *IFNB1*, or (c) *ISG15* or (d) whole cell lysates were analysed by western blot analysis for viral NP, IFIT1 and or ϕ3-Actin protein expression. Poly(I:C) transfected A549 cells (24 h post-transfection) were used as control where indicated. (e-f) A549 cells were infected with (d) MOBV or (e) LASV at an MOI of 0.1. At 48 hpi, cells were fixed and analysed by immunofluorescence staining for viral NP protein and DAPI staining for DNA.

**Extended Data Figure 2:**
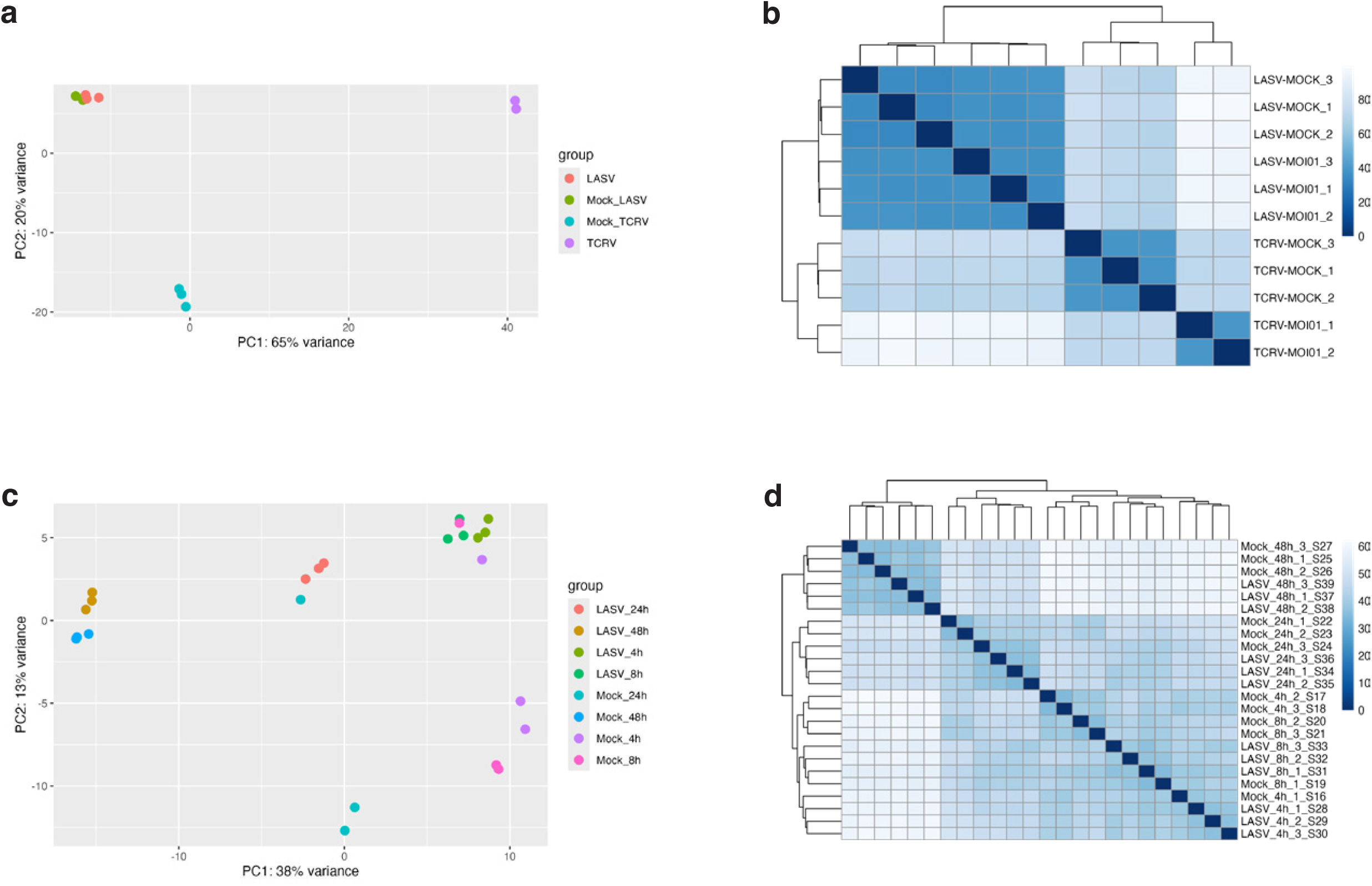
TCRV but not LASV induces comprehensive transcriptional changes in A549 cells. (a-b) A549 cells were infected with TCRV or LASV at an MOI of 0.1 for 48 hpi and total RNA was analysed by total RNA-seq. Individual samples were globally compared based on their gene expression profiles using (a) principal component analysis (PCA) and a (b) distance matrix. (c-d) A549 cells were infected with LASV at an MOI of 3 and total RNA was analysed by total RNA-seq at 4 to 48 hpi. Individual samples were globally compared based on their gene expression profiles using (c) principal component analysis (PCA) and a (d) distance matrix.

**Extended Data Figure 3:**
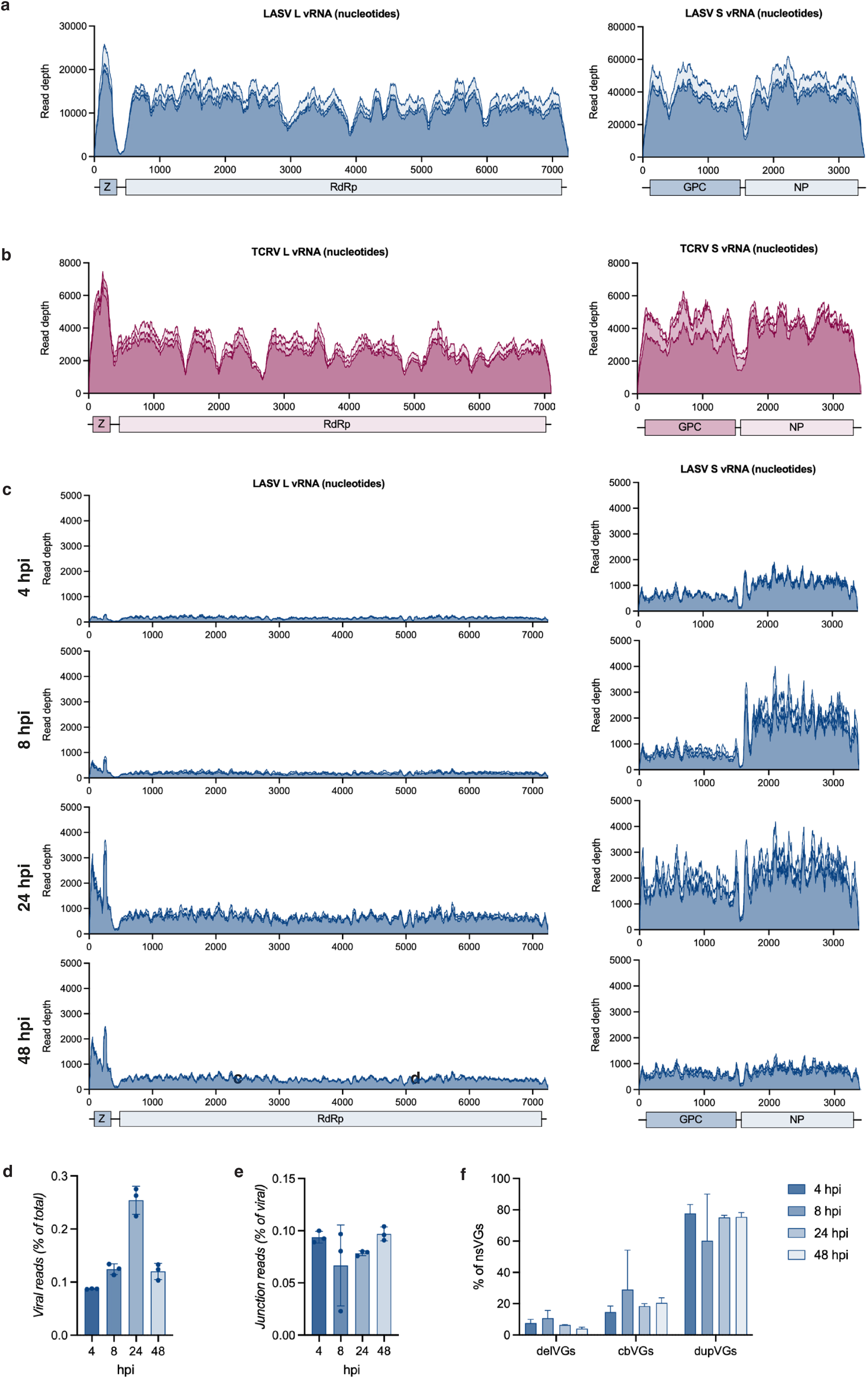
Analysis of TCRV and LASV nsVGs produced in A549 cells. (a-b) A549 cells were infected with LASV or TCRV (MOI = 0.1) for 48 hpi and total RNA was analysed by total RNA-seq. The graphs depict the per nucleotide read depth along the (a) LASV or (b) TCRV genome from three independent biological replicates after bowtie2 alignment. (c-f) A549 cells were infected with LASV (MOI = 3) for 4 to 48 hpi and total RNA was analysed by total RNA-seq. (c) The graphs depict the per nucleotide read depth along the LASV genome from three independent biological replicates after bowtie2 alignment. (d) Relative continuous alignment rate using bowtie2 to the LASV reference genome compared to total reads. (e) Relative discontinuous alignment rate using ViReMa to the LASV reference genome compared to total continuous viral reads.(f) Relative frequency of different nsVG types.

**Extended Data Figure 4:**
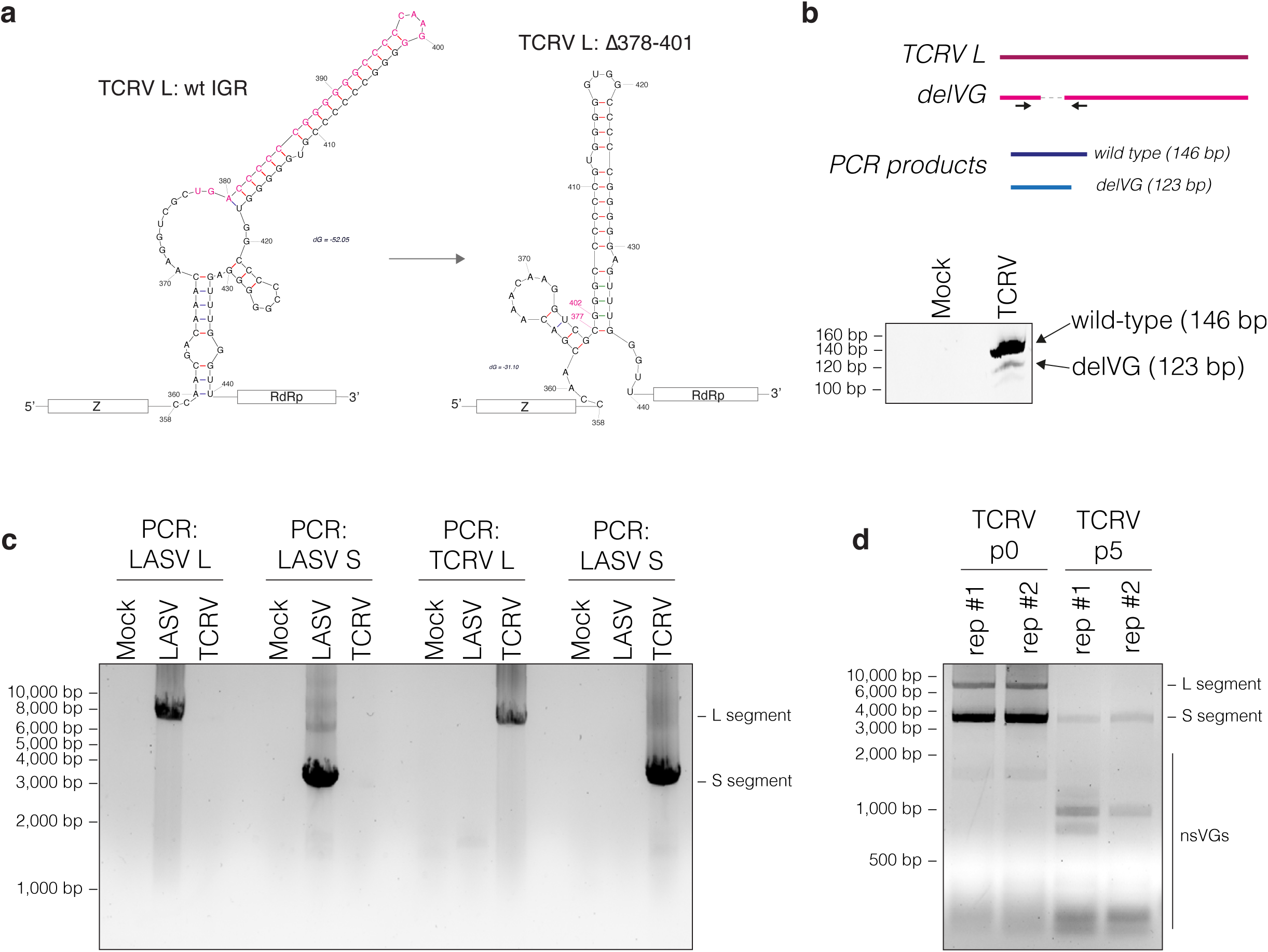
Validation of arenavirus nsVG production. (a) Schematic of the secondary structure of the TCRV L segment IGR with or without the identified delVG truncation Δ378-401. The deleted sequence and the junction coordinates are highlighted in red. (b) RT-PCR to validate the presence of TCRV L segment IGR truncations. PCR products were analysed by denaturing 7M urea 10% PAGE. (c) RNA from A549 cells infected with LASV or TCRV (MOI = 0.1) for 48 hpi was analysed by vgRT-PCR. (d) TCRV was blindly passaged for 5 passages in Vero E6 cells. RNA from Vero E6 cells infected with unpassaged and passaged TCRV was analysed by vgRT-PCR.

